# Role of FBXW2 in explant culture of bovine periosteum-derived cells

**DOI:** 10.1101/2020.12.17.423216

**Authors:** Mari Akiyma

**Affiliations:** Department of Biomaterials, Osaka Dental University, 8-1, Kuzuhahanozono-cho, Hirakata-shi, Osaka 573-1121, Japan

**Author notes:** Corresponding author (MA).

## Abstract

Osteoporosis and bone fracture decrease quality of life. Bone regeneration is a notable technique for osteoporosis treatment. A previous study reported that F-box and WD-40 domain-containing protein 2 (FBXW2) and osteocalcin have the same shape in the periosteum after 5 weeks. However, the osteoblastic functions of FBXW2 are not clear. In this study, double fluorescent immunostaining revealed a small amount of osteocalcin in the area of FBXW2 aggregation at 1 week, periosteal cells, and osteocalcin pushed toward the edge of periosteum, and, apart from FBXW2 tubes at 2 weeks, multilayered periosteum-derived cells at 3 weeks and sticking of osteocalcin in the periosteum with cells at 4 weeks. At 5 weeks, FBXW2 disappeared at the root of periosteum-derived cells, while osteocalcin and cells remained. Based on these results, it is hypothesized that FBXW2 maintains tissue shapes and prevents escape of inner periosteal cells, and the disappearance of FBXW2 causes migration of periosteum-derived cells out of the periosteum along with osteocalcin. Furthermore, FBXW2 may play a role in dynamic tissue remodeling and bone formation.

## Introduction

Osteoporosis and bone fracture decrease quality of life. Bisphosphonates used for the treatment of osteoporosis are associated with the risk of osteonecrosis of the jaw [1]. Besides bisphosphonates, bone regeneration is a notable treatment for osteoporosis. Many studies have investigated the role of the cambium layer of the periosteum [2–5]; however, specific proteins in the periosteum that aid in bone formation are still unknown. Periosteal stem cells are also important for bone regeneration [6]. Bovine periosteum-derived cells are used for bone regeneration, and these cells can form multilayered cell sheets without scaffolds on tissue culture dishes [7]. To determine the mechanism of multilayered cell sheet formation, the supernatant and periosteum were studied [8,9]. Akiyama investigated the supernatant of bovine periosteum-derived cells using mass spectrometry and immunohistochemistry [9] and found that F-box and WD-40 domain-containing protein 2 (FBXW2) is expressed in the periosteum [10]. FBXW2 is one of the F-box proteins involved in the ubiquitin-proteasome system [11]. Among the 69 known F-box proteins, only four—FBXW7, SKP2, β-TRCP1, and β-TRCP2—are well studied, whereas the functions of the remaining 65 members are still unknown [12]. Thus, the function of FBXW2 is also unknown. In 2018, Akiyama reported that FBXW2 and osteocalcin form tubes of the same shape in the periosteum after 5 weeks in culture [13], but the relationship between these two proteins is not clear. The osteoblastic function of FBXW2 is also unknown. In this study, periosteum and periosteum-derived cells were observed for up to 5 weeks using double fluorescent immunostaining to determine the effects of FBXW2 on the osteoblastic character of periosteum-derived cells.

## Materials and methods

### Preparation of periosteum

All protocols were approved by the Animal Research Committee of Osaka Dental University (approval number 20–02006) and complied with fundamental guidelines for proper conduct of animal experiments and related activities in academic research institutions under the jurisdiction of the Ministry of Education, Culture, Sports, Science and Technology (The Ministry of Education, Culture, Sports, Science and Technology directive 2006, Notice No. 71). The periosteum was separated from the bovine leg (Kobe Chuo Chikusan, Kobe, Japan) as described previously [7]. Bone sections were removed, fixed with 4% paraformaldehyde (PFA), and cast into paraffin blocks. The periosteum was cultured in 100 mm dishes in Medium 199 supplemented with 10% fetal bovine serum, 100 units penicillin, 100 μg streptomycin/mL solution (Wako Pure Chemical Industries, Ltd., Osaka, Japan), and 5 mg/mL ascorbic acid for up to 5 weeks. The medium was changed once a week. The periosteum was fixed every week and prepared for sections. Figure 1 shows the schema of this study.

**Fig 1.**
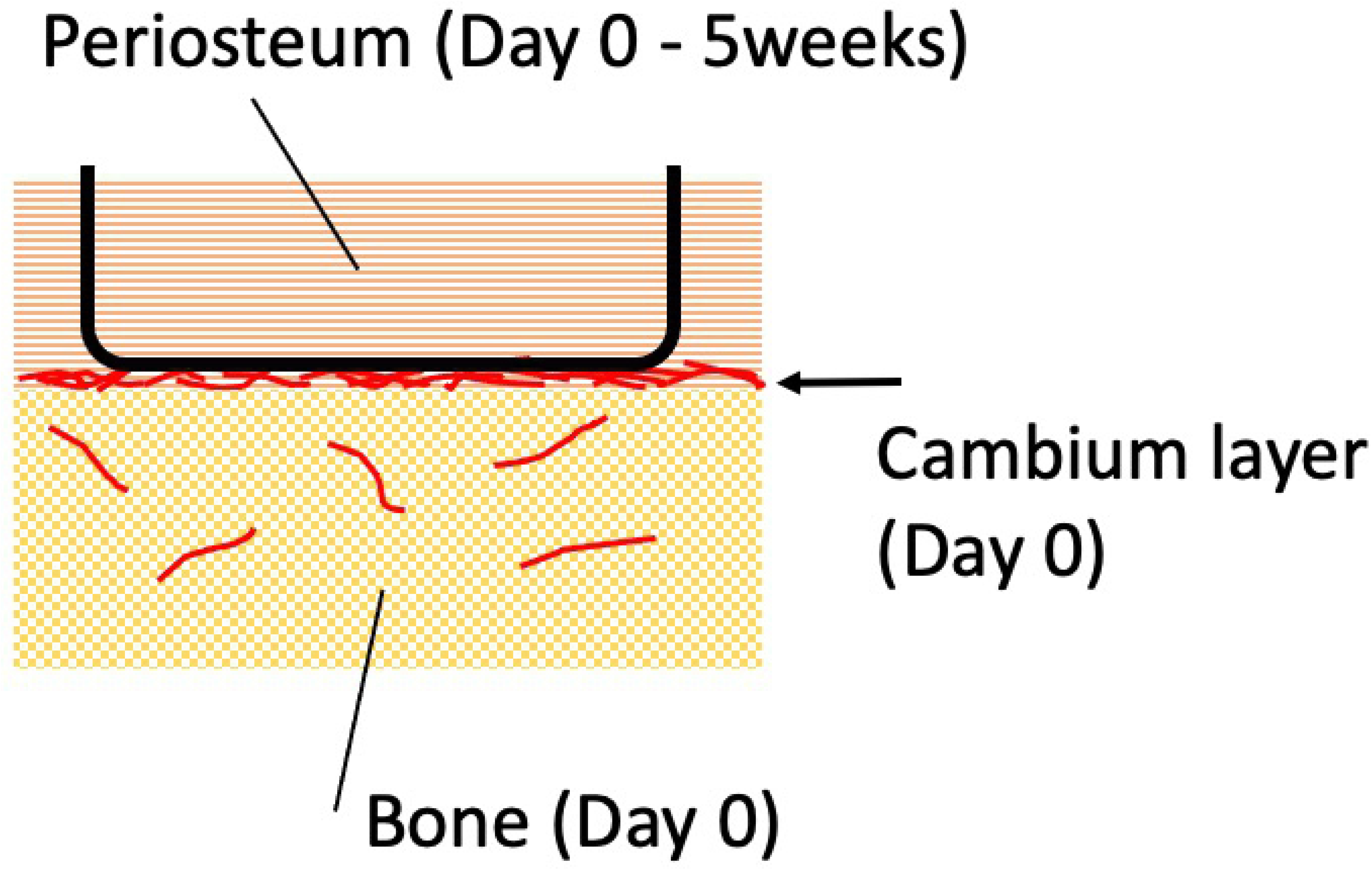
Schema of this study.

### Fluorescent immunostaining and immunohistochemistry

All paraffin sections were pre-treated with ready-to-use Proteinase K (Dako Cytomation, Glostrup, Denmark) for 10 min. Primary antibodies used were mouse monoclonal antibodies for osteocalcin (Santa Cruz Biotechnology, Inc., No. sc-376835), a mouse monoclonal antibody for bovine osteocalcin (code no. M042, clone no. OCG2; Takara Bio Inc., Shiga, Japan), and goat polyclonal antibody for FBXW2 (Invitrogen, #PA5-18189). The secondary antibodies used were Alexa Fluor™ 488 goat anti-mouse (Invitrogen, #A11029), mouse anti-goat IgG-CFL 594 (Santa Cruz Biotechnology, Inc., No.sc516243), and N-Histofine Simple Stain AP (multi) (#414261, Nichirei Biosciences Inc., Tokyo, Japan). Alkaline phosphatase-tagged antibody was visualized with PermaRed/AP (K049, Diagnostic BioSystems, CA, USA). Incubation with mouse monoclonal antibodies for osteocalcin (diluted 1:100) was performed overnight at 4 °C, and that with monoclonal antibody for bovine osteocalcin (diluted 1:500) and FBXW2 antibody (diluted 1:100) was performed for 4 h at room temperature. Cell nuclei were stained with hematoxylin or DAPI. For negative controls, the antibody for receptor activator of NF-κB ligand (RANKL) (Santa Cruz Biotechnology, Inc., No. sc-377079) and normal goat serum (Fuji film Wako Pure Chemical Industries, Ltd. 143–06561) were used. Images were photographed with a fluorescence microscope (Keyence Japan, Osaka, Japan, BZ-9000).

## Results

Figure 2 shows the bone and cambium layer. Consistent with a previous study [13], FBXW2 (red) is expressed in the bone and cambium layer (Fig 2(a)). Figure 2(b) shows negative control of the anti-FBXW2 goat antibody (red); only cell nuclei (blue) are stained. Figure 2(c) shows osteocalcin expression using a mouse monoclonal antibody for osteocalcin (No. sc-376835). As shown in Fig 2(d), RANKL is not expressed. An alkaline phosphatase labeled secondary antibody was used in Fig 2(c)–(d), and fluorescent labeled secondary antibody was used in Fig 2(e)–(f). In Fig 2(c)–(e), the same primary antibody for osteocalcin was used. Figure 2(e) reveals that osteocalcin is expressed in bone (green), but not in the cambium layer (cell nuclei stained blue). In Fig 2(d) and 2(f), the same primary antibody for RANKL was used. Figure 3 shows double fluorescent immunostaining of the periosteum at day 0. In Fig 3, FBXW2 is expressed in blood vessels, while osteocalcin is not. Although osteocalcin reacts with blood cells, the reaction may be non-specific.

**Fig 2.**
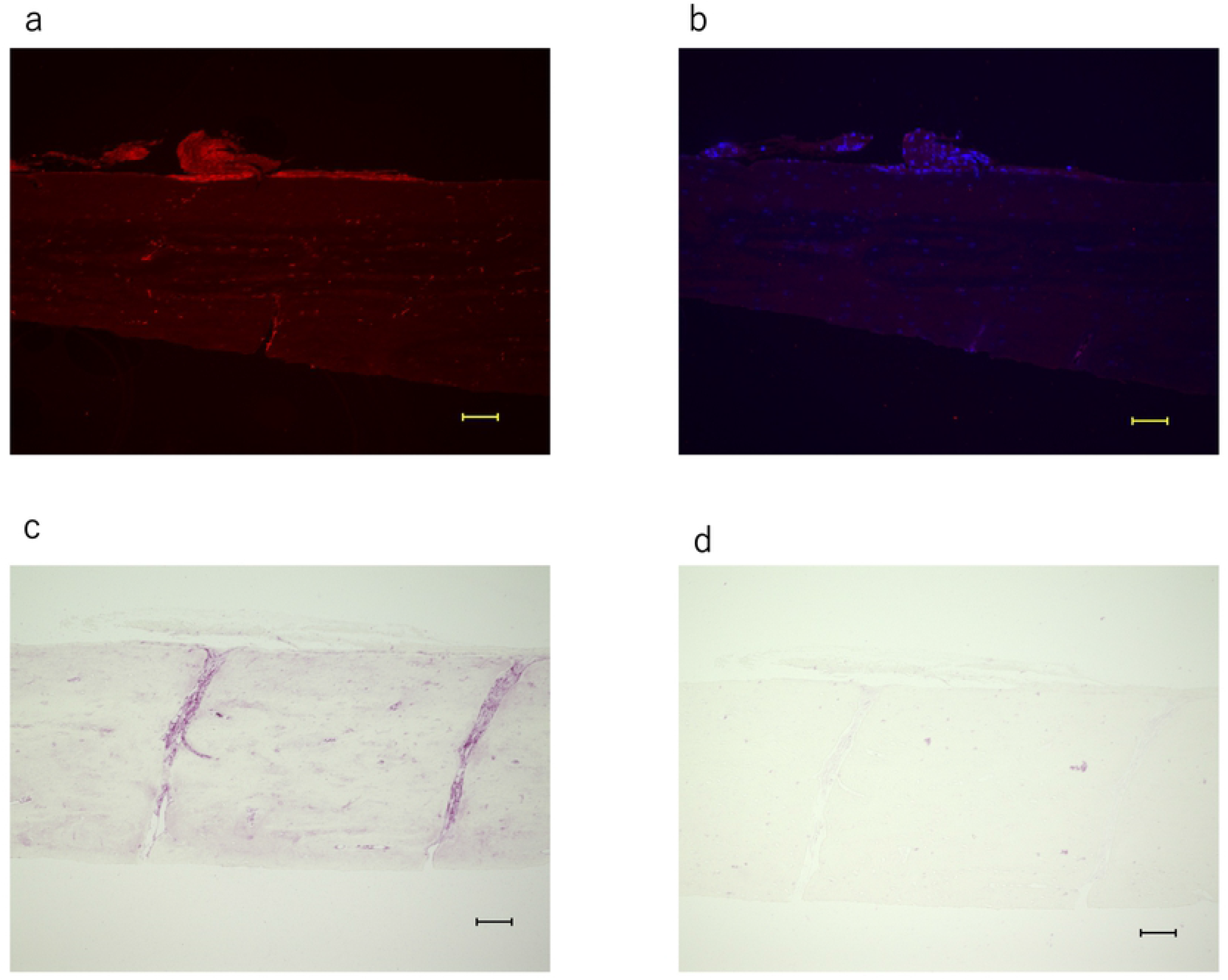
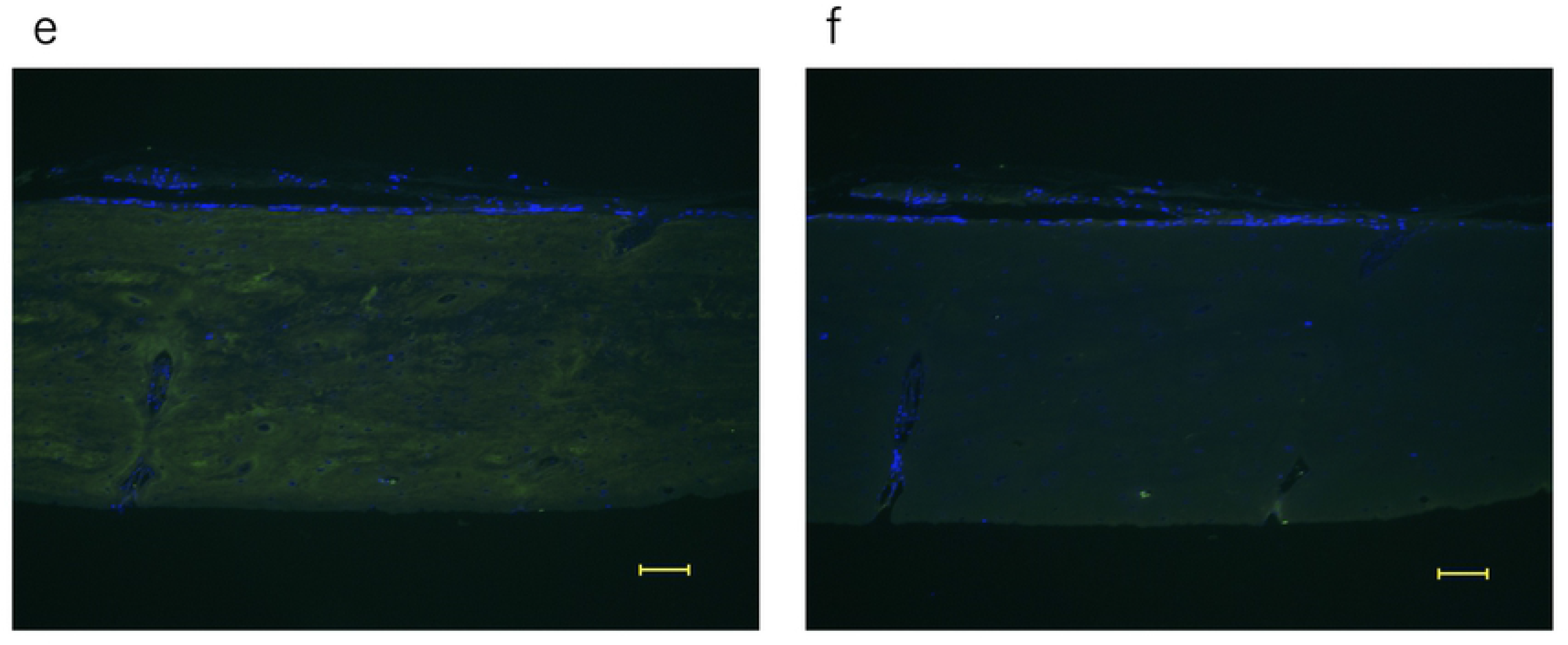
Fluorescent immunostaining and immunohistochemistry of bone. Scale bar: 100μm. (a) FBXW2: red, (b)negative control of (a), (c) osteocalcin, (d) RANKL: negative control of (c), (e) osteocalcin: green, (f) RANKL: negative control of (e)

**Fig 3.**
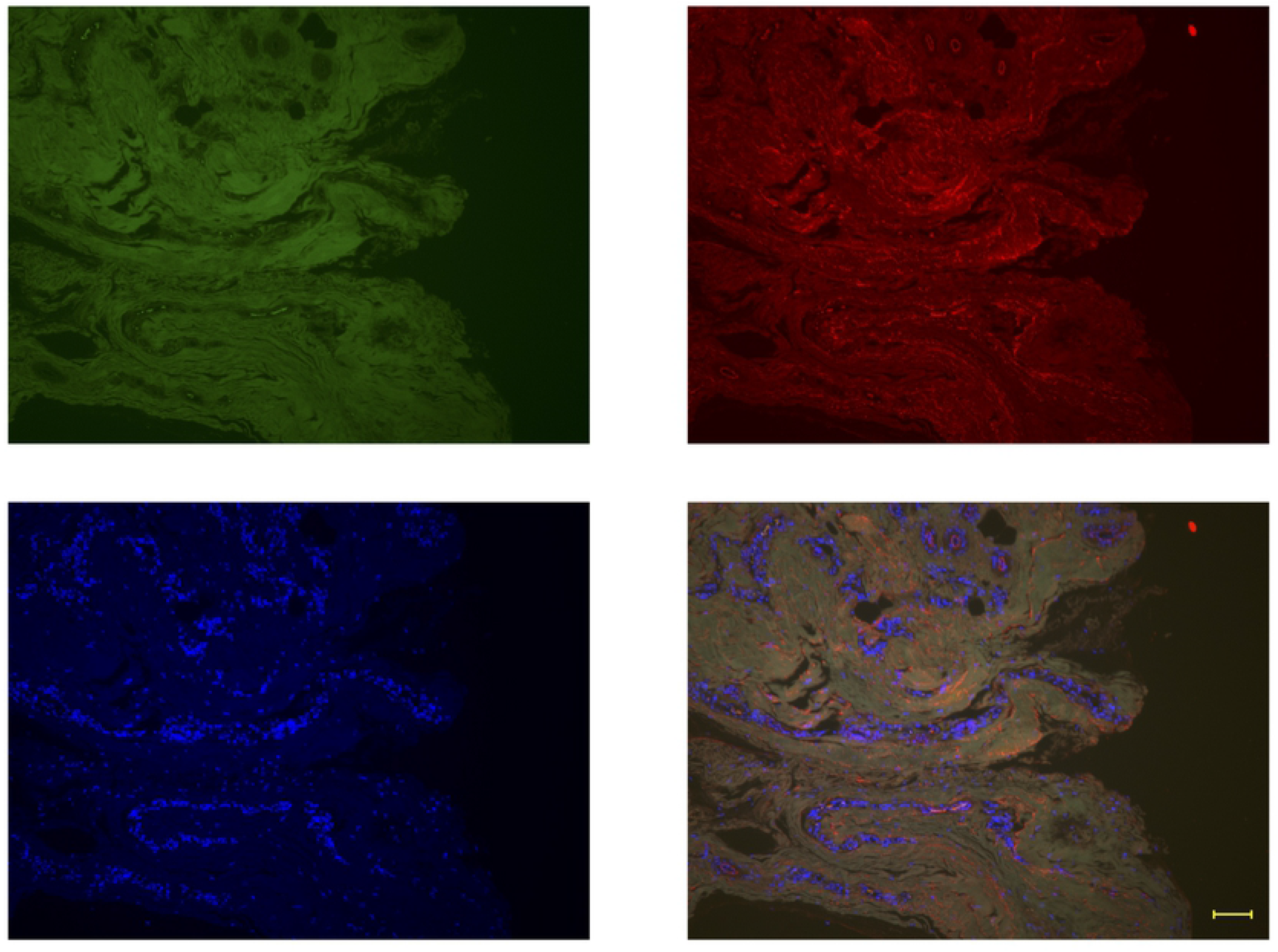
Double fluorescent immunostaining of periosteum at day 0. Osteocalcin: green, FBXW2: red, DAPI: blue Scale bar: 100μm.

Monoclonal antibody for bovine osteocalcin (code no. M042) was used for double fluorescent immunostaining at week 1 and continued for up to 5 weeks. Figure 4(a) shows that osteocalcin is expressed near FBXW2, but it is localized to small regions. Figure 4(b) shows blood vessels at 1 week and FBXW2 expression in blood vessels. Figure 4(c)–(f) show high magnification of bovine periosteal cells, which may synthesize osteocalcin. Shape of osteocalcin-resembled cells (Fig 4(c),(d)), stick (Fig 4(e)), and FBXW2 (Fig 4(f)). Figure 5a-e show the expression of osteocalcin along multiple edges of the periosteum; expression of osteocalcin increased with respect to observations at 1 week. At 2 weeks, FBXW2 expression decreased along the edges of periosteum, while cells and osteocalcin remained the same (Fig 5(a)–(e)). Expression of RANKL, which was used as a negative control, was absent (Fig 5(f)). Osteocalcin expression along the edges increased at 3 weeks (Fig 6(a)) and multiple layers of periosteum-derived cells appeared (Fig 6(b)). Osteocalcin appeared in the regions where FBXW2 levels had decreased (Fig 6(c)). At 4 weeks, a stick of osteocalcin poked out of the periosteum with multiple layers of periosteum-derived cells (Fig 7). At 5 weeks, FBXW2 disappeared from the regions where periosteum-derived cells had appeared (Fig 8(a),(b)), while cells and osteocalcin remained unaffected (Fig 8(b)–(d)).

**Fig 4.**
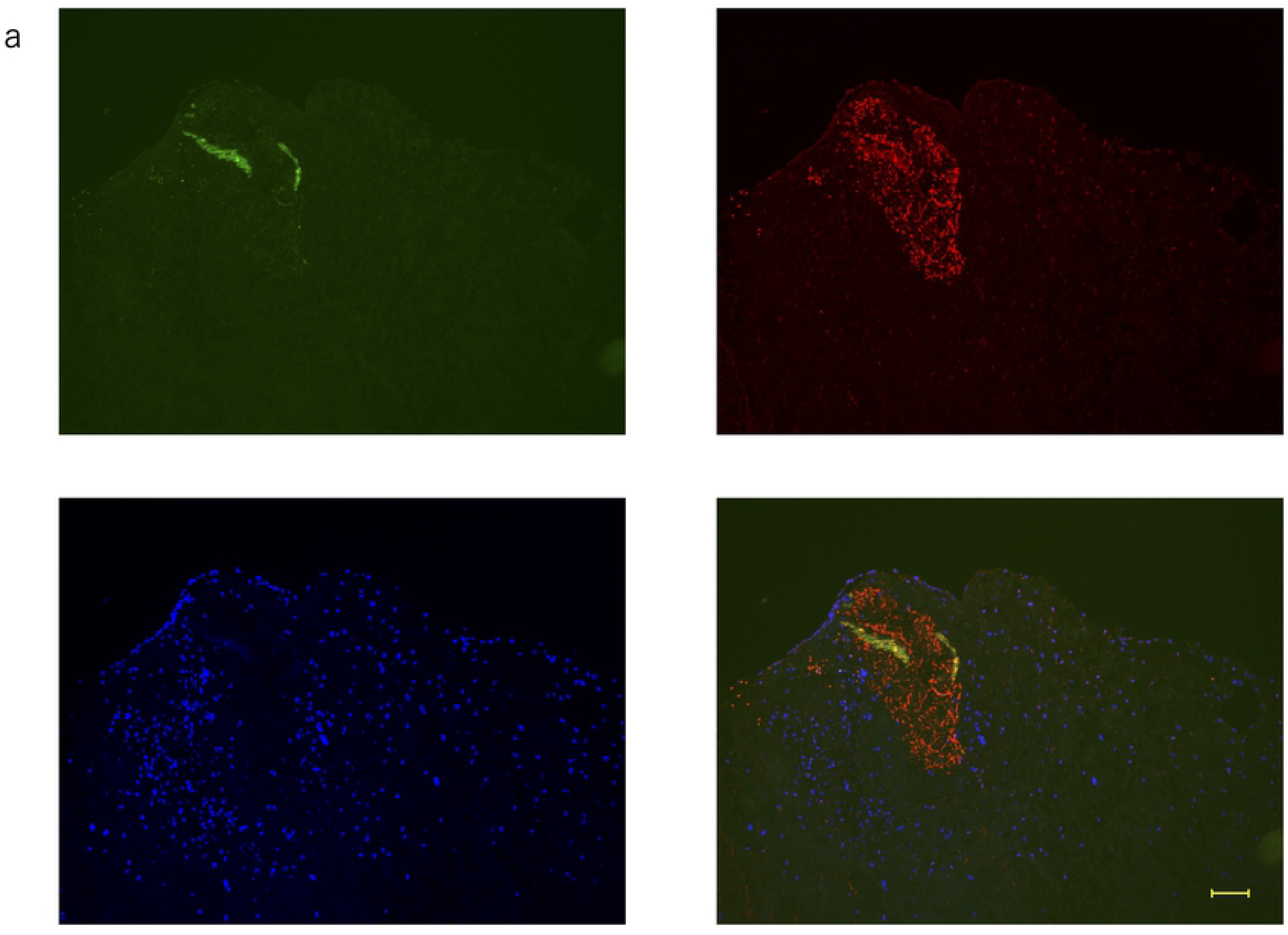
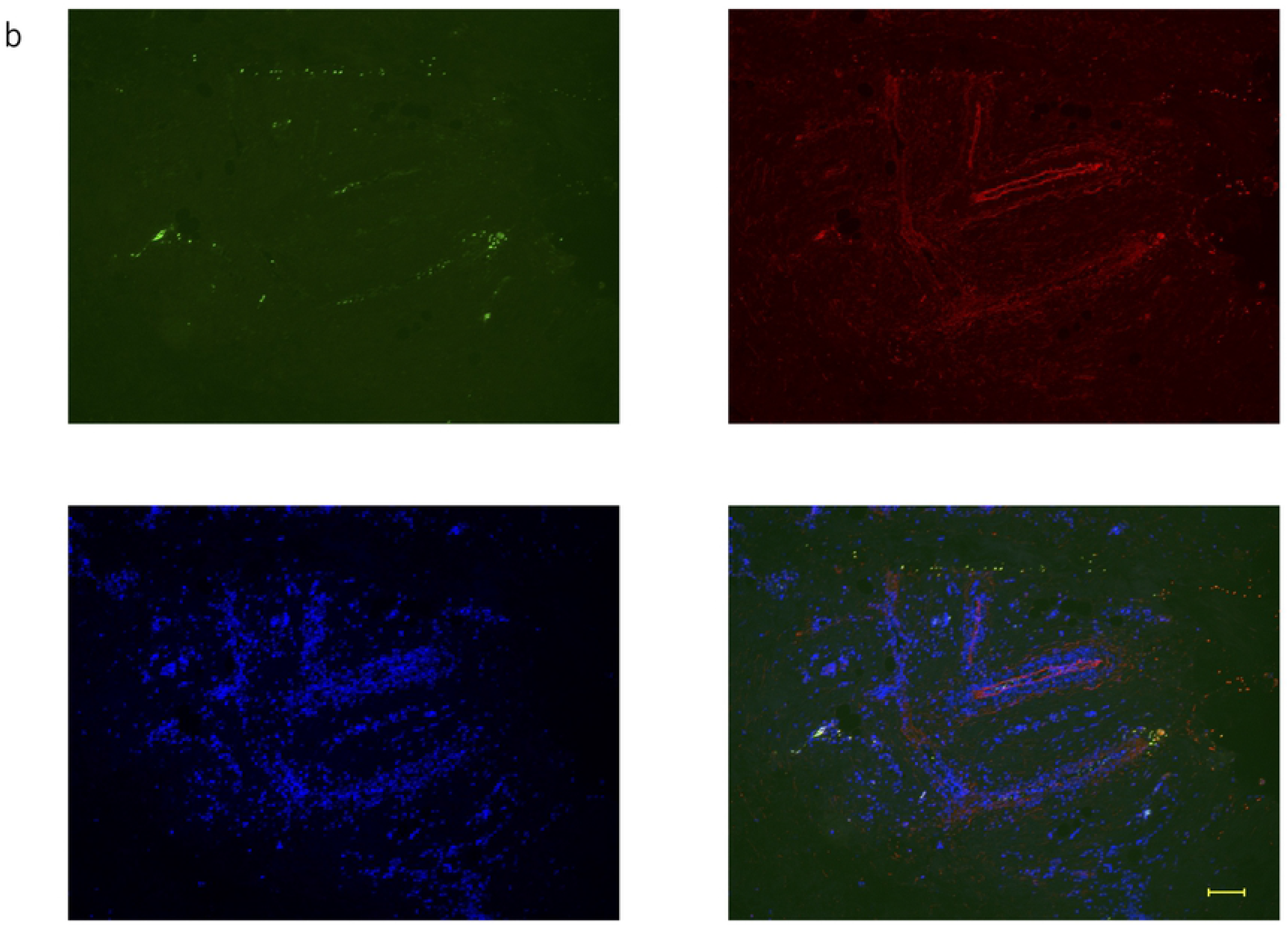
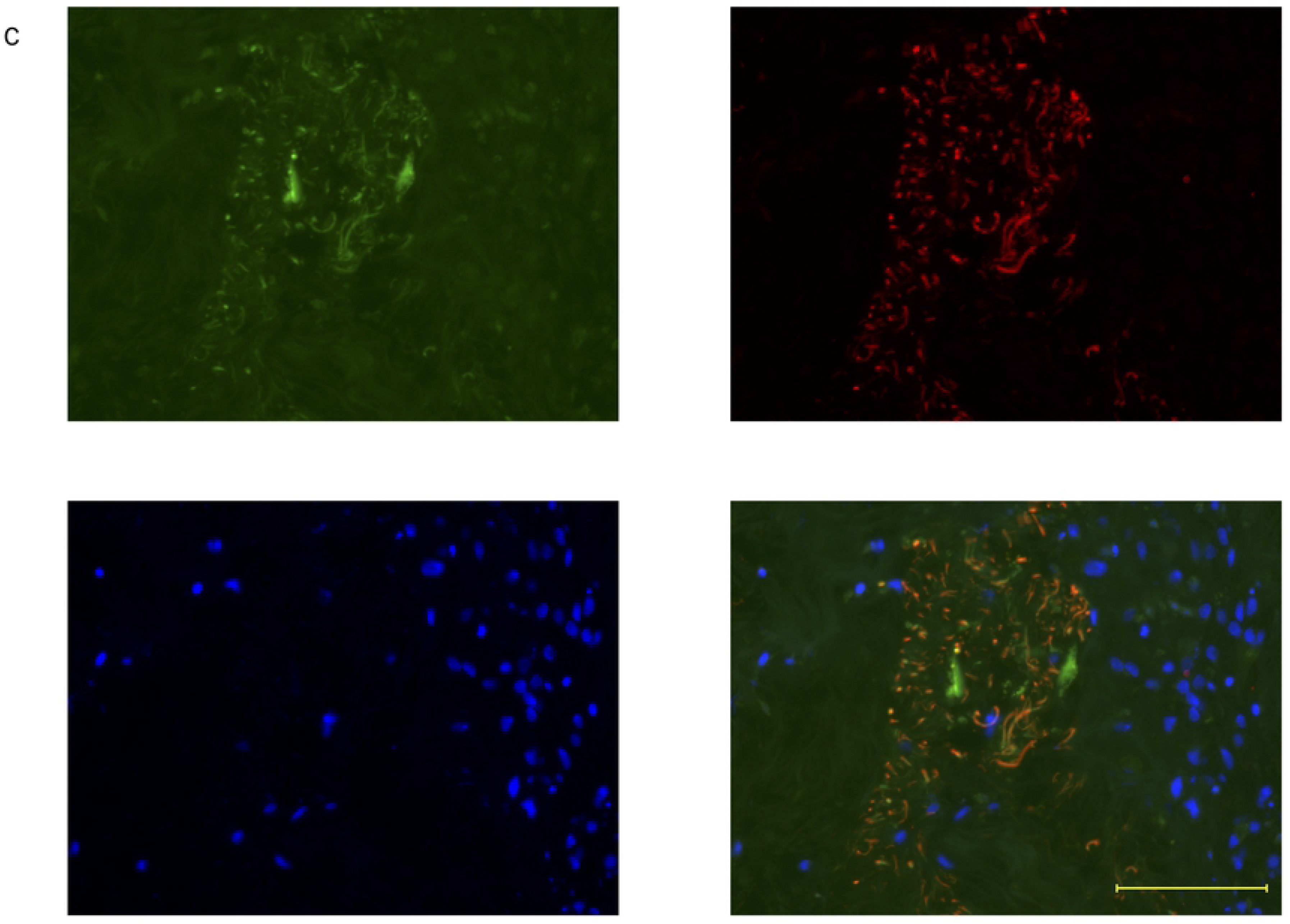
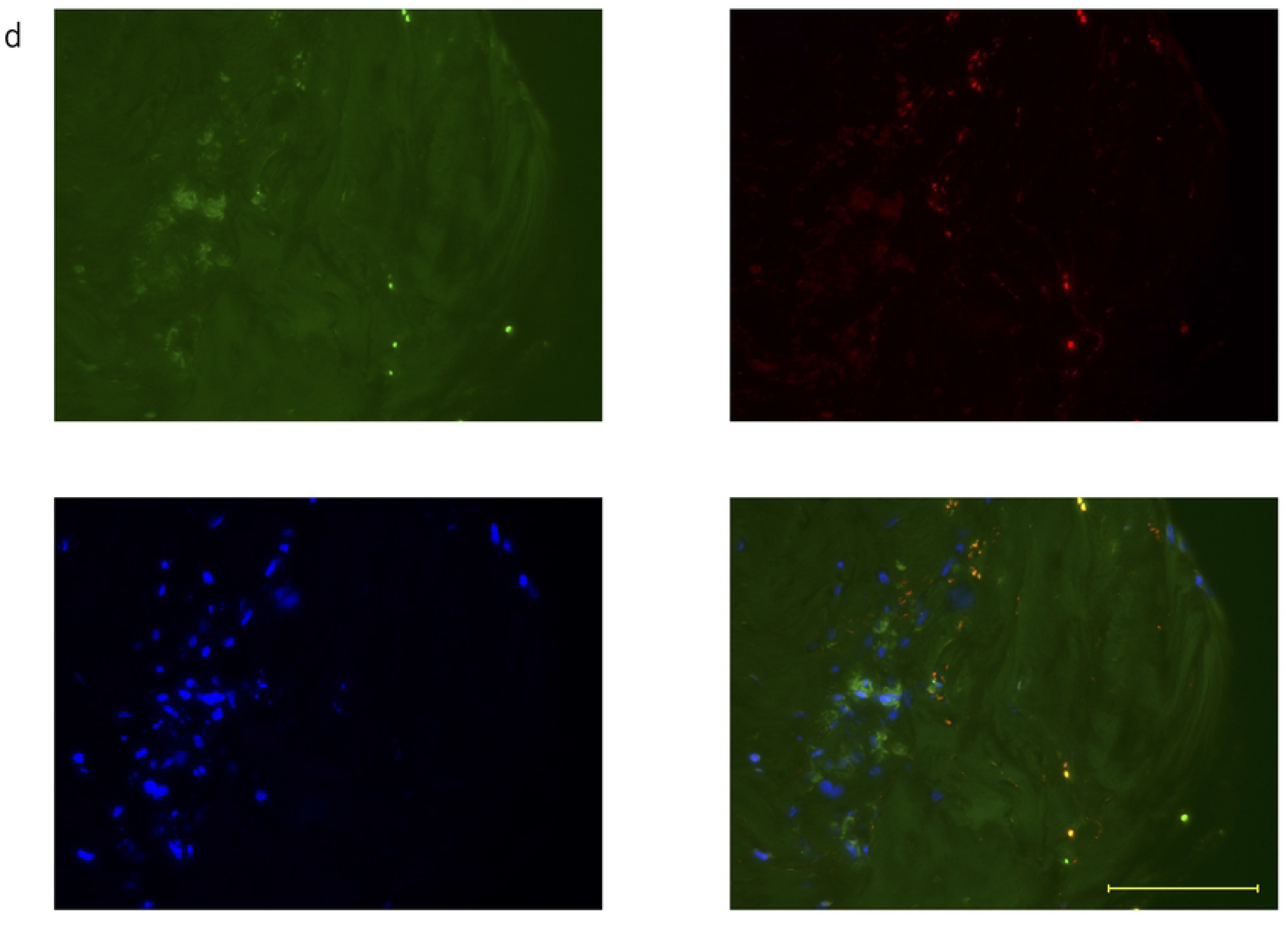
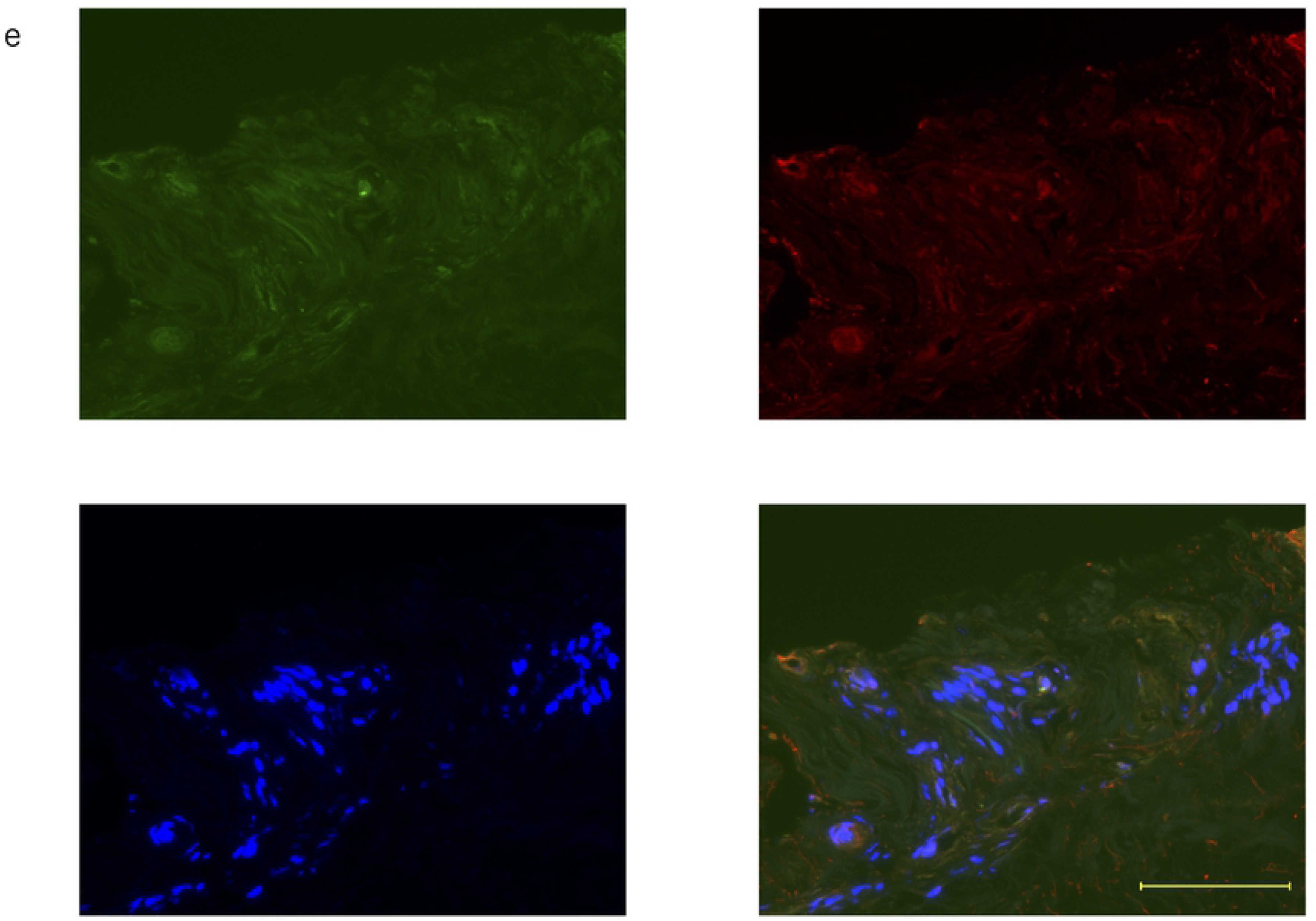
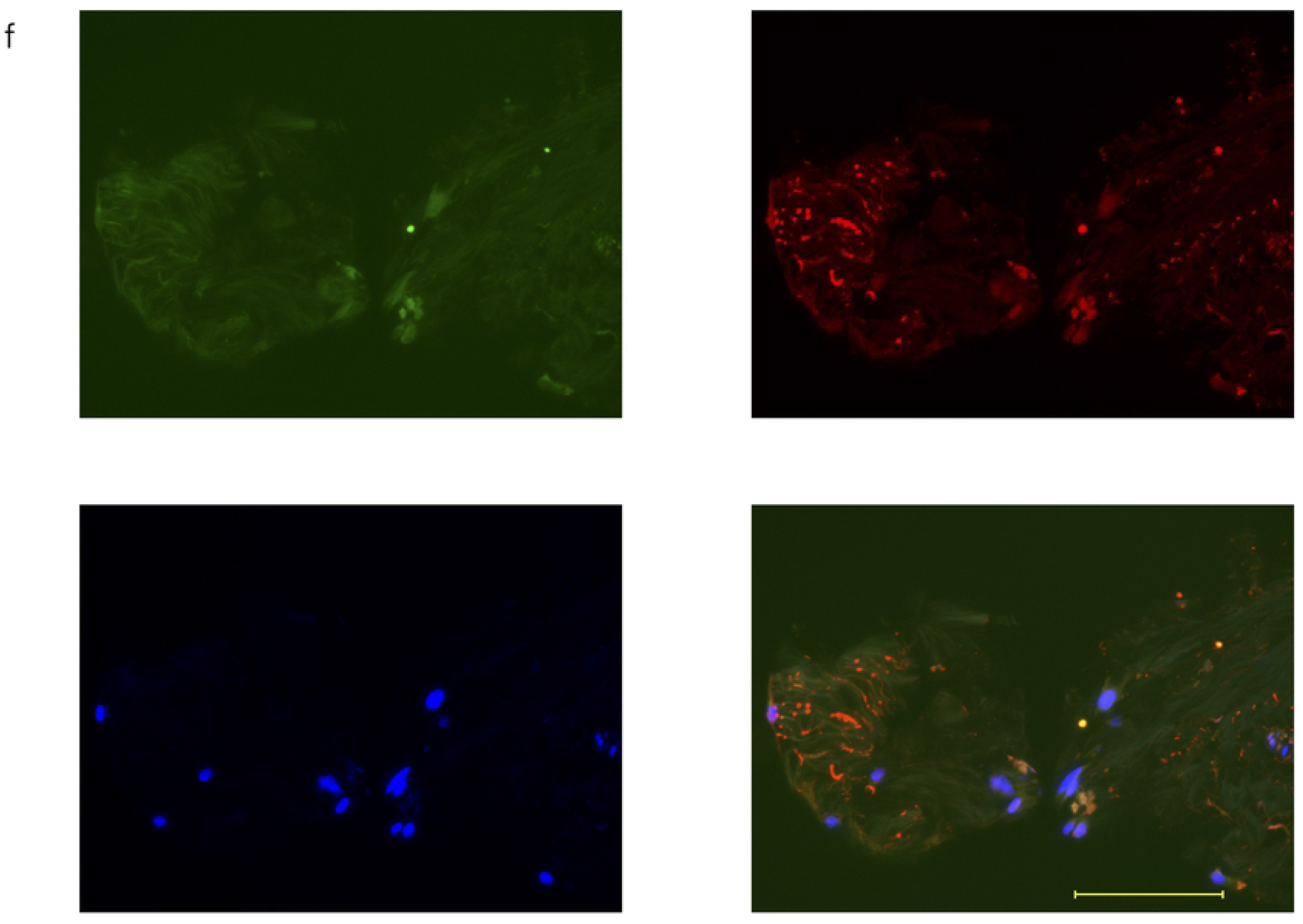
(a)-(f) Double fluorescent immunostaining of periosteum at 1 week. Scale bar: 100μm. Osteocalcin: green, FBXW2: red, DAPI: blue

**Fig 5.**
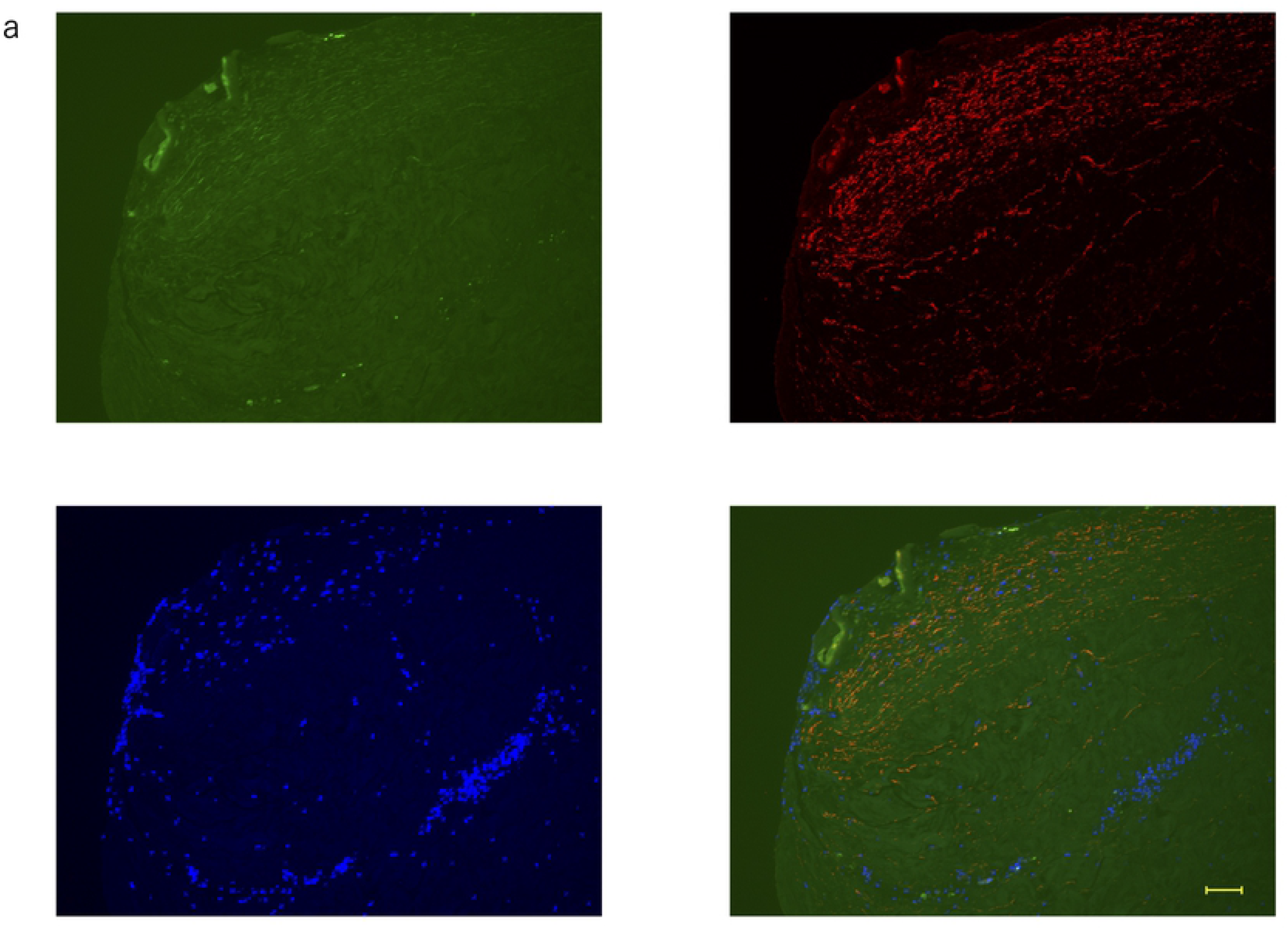
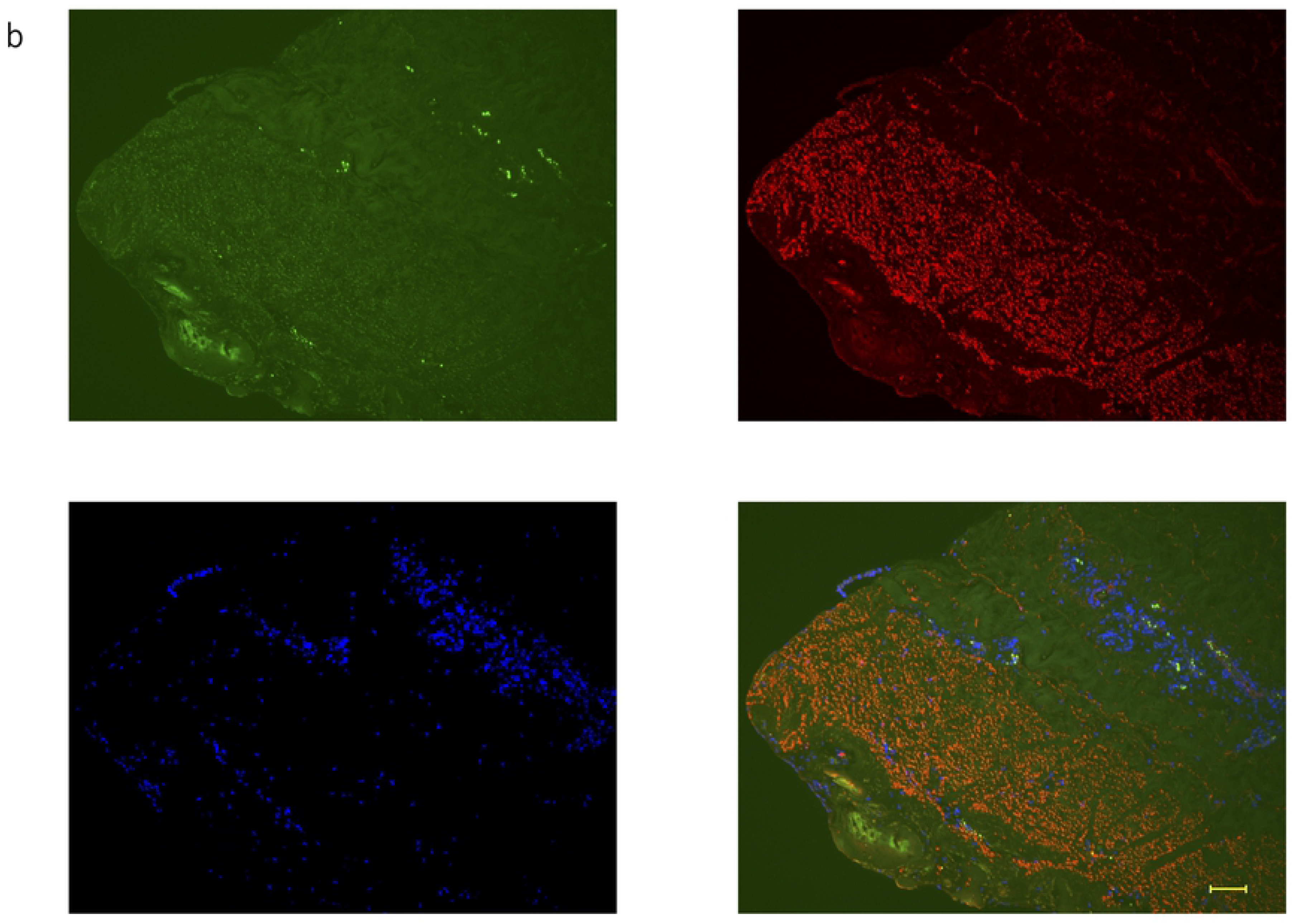
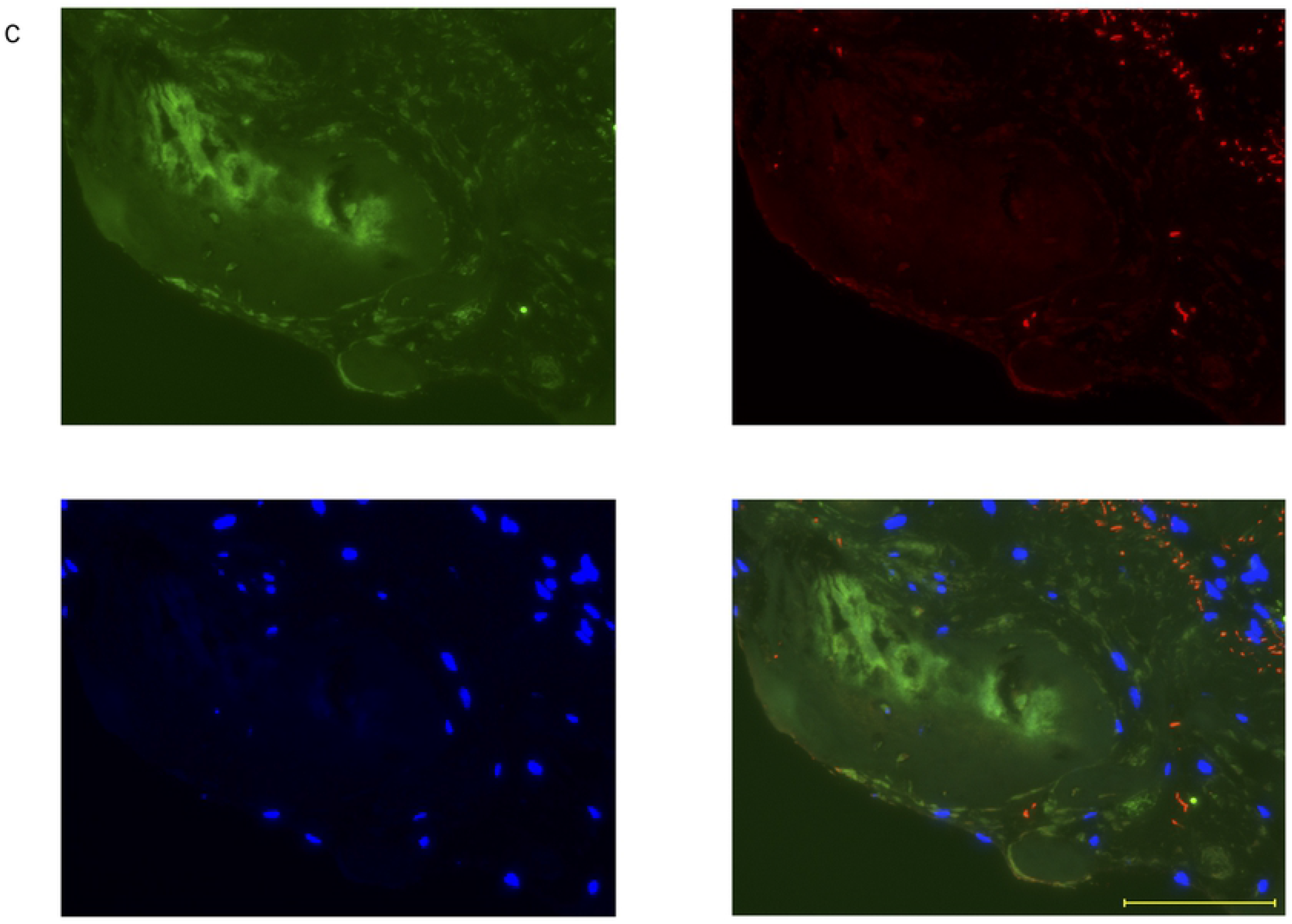
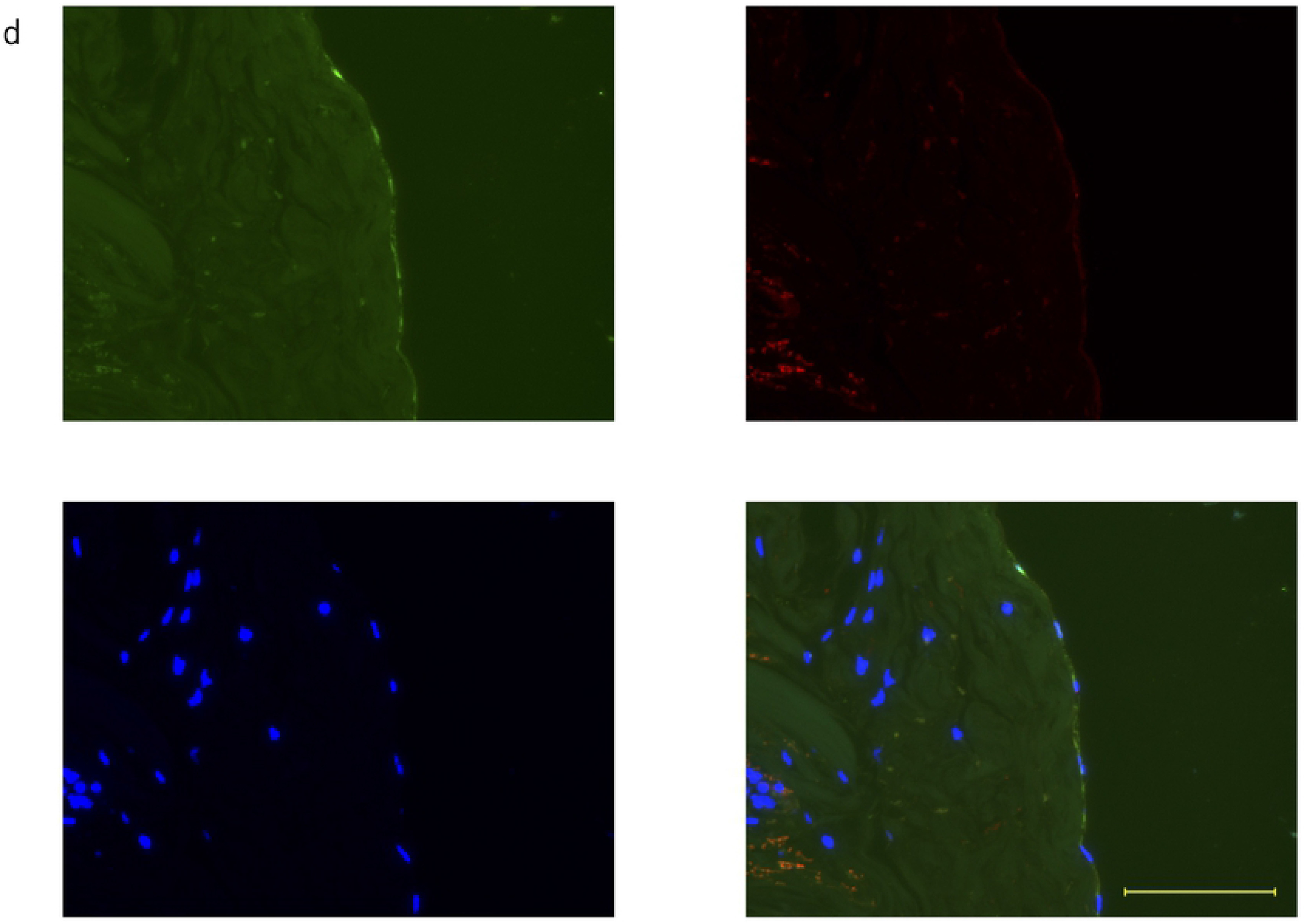
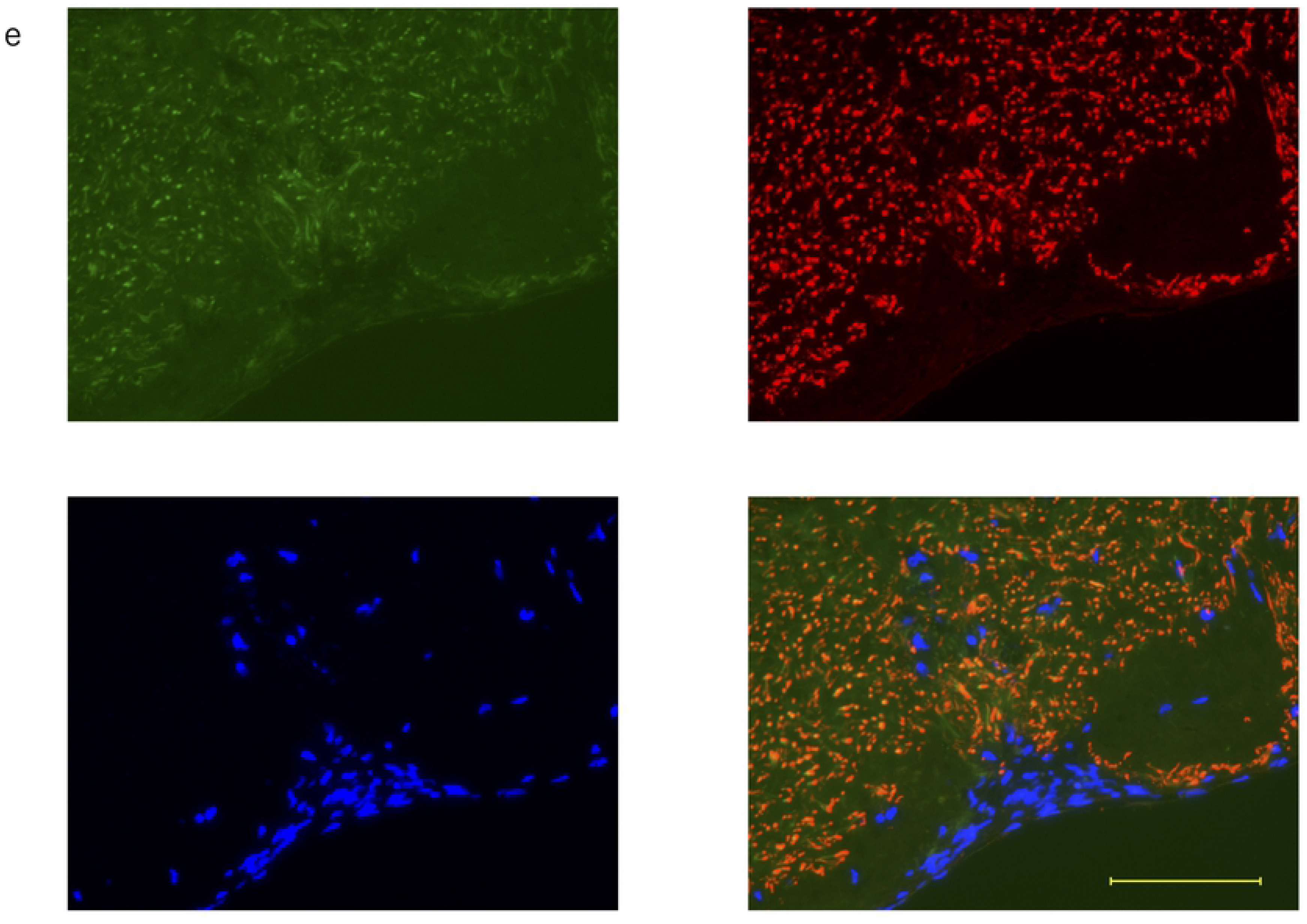
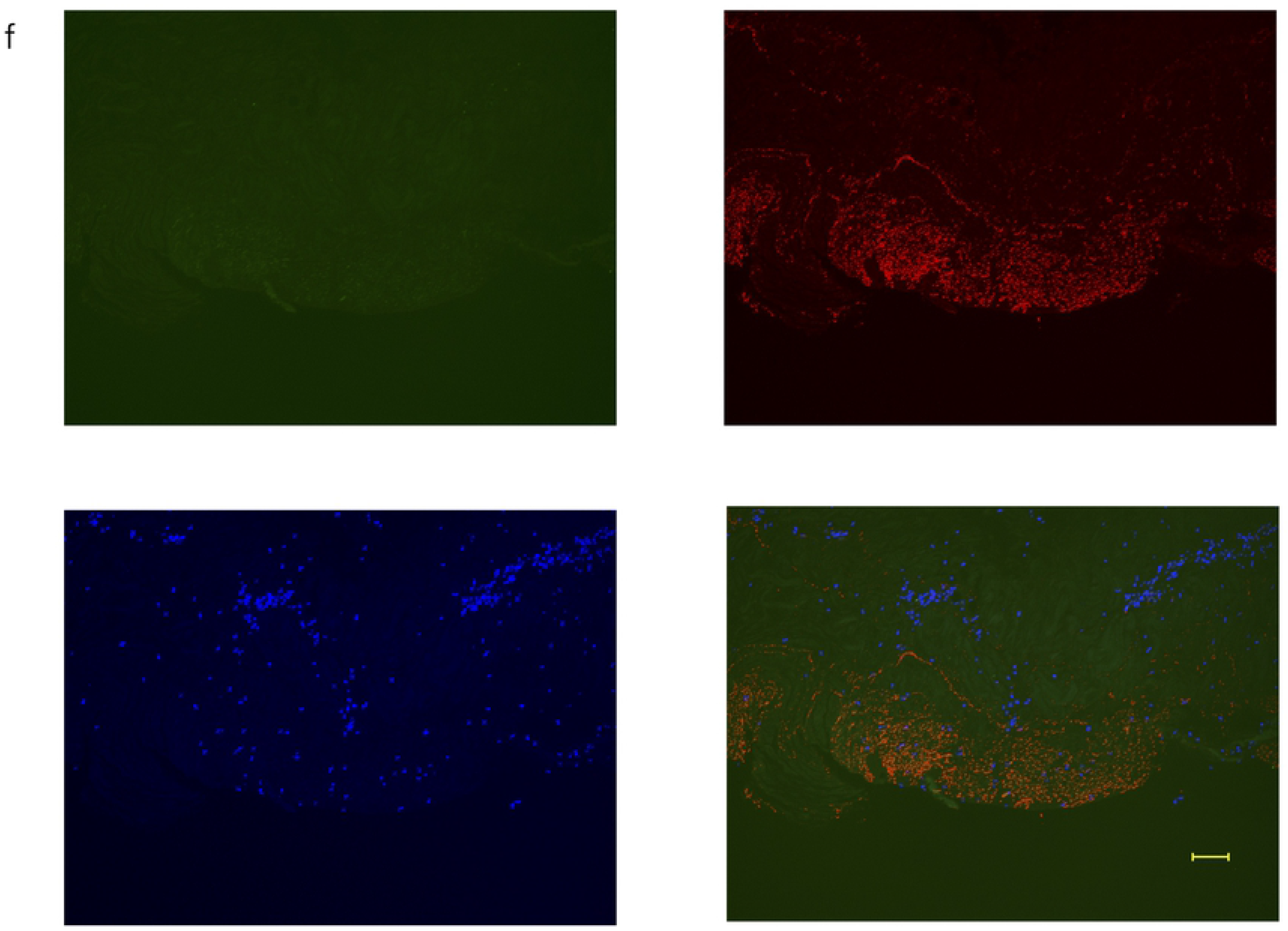
(a)-(e) Double fluorescent immunostaining of periosteum at 2 weeks. Scale bar: 100 μm. Osteocalcin: green, FBXW2: red, DAPI: blue; (f) negative control, RANKL: green

**Fig 6.**
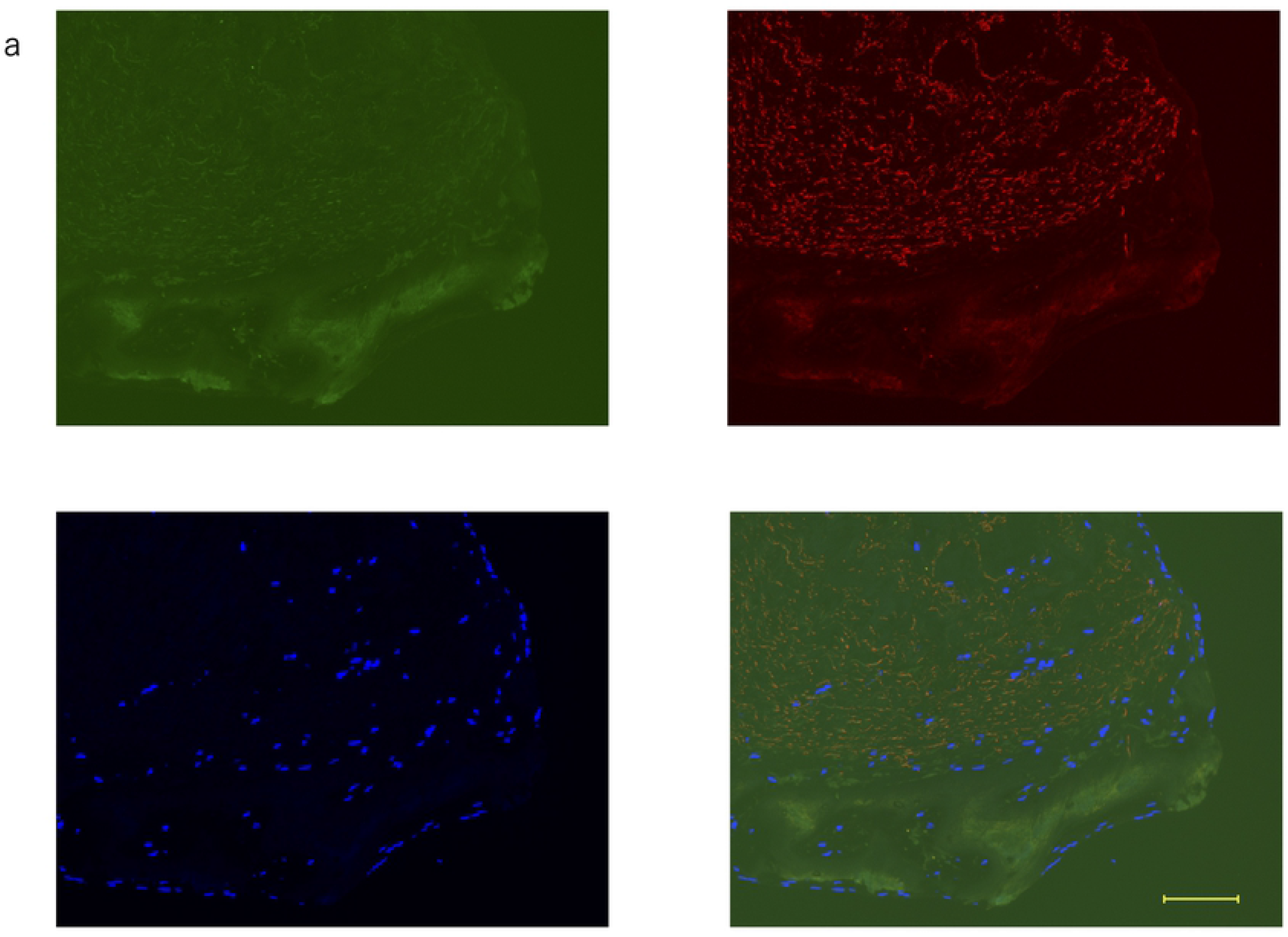
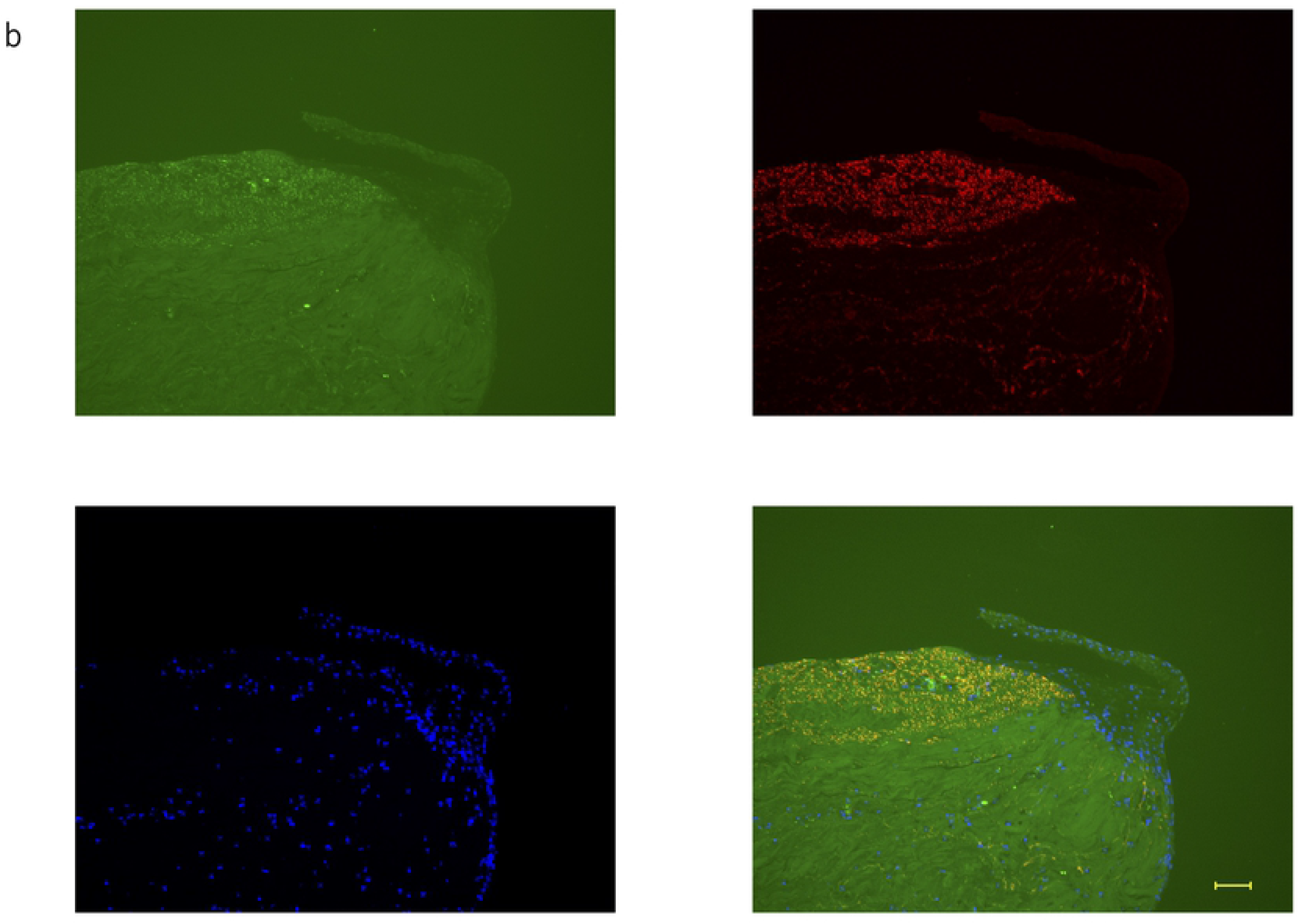
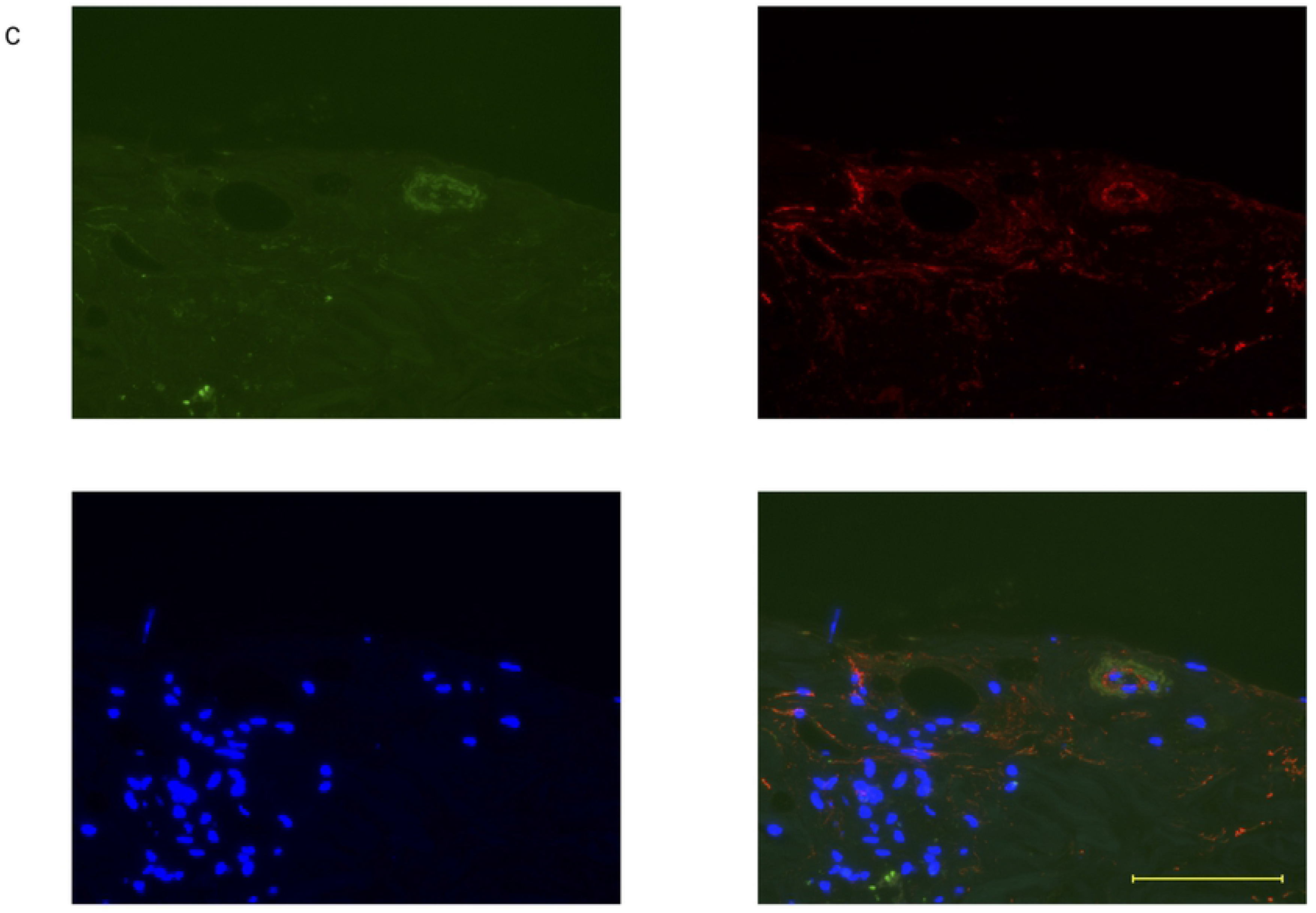
(a)-(c) Double fluorescent immunostaining of periosteum at 3 weeks. Scale bar: 100 μm. Osteocalcin: green, FBXW2: red, DAPI: blue

**Fig 7.**
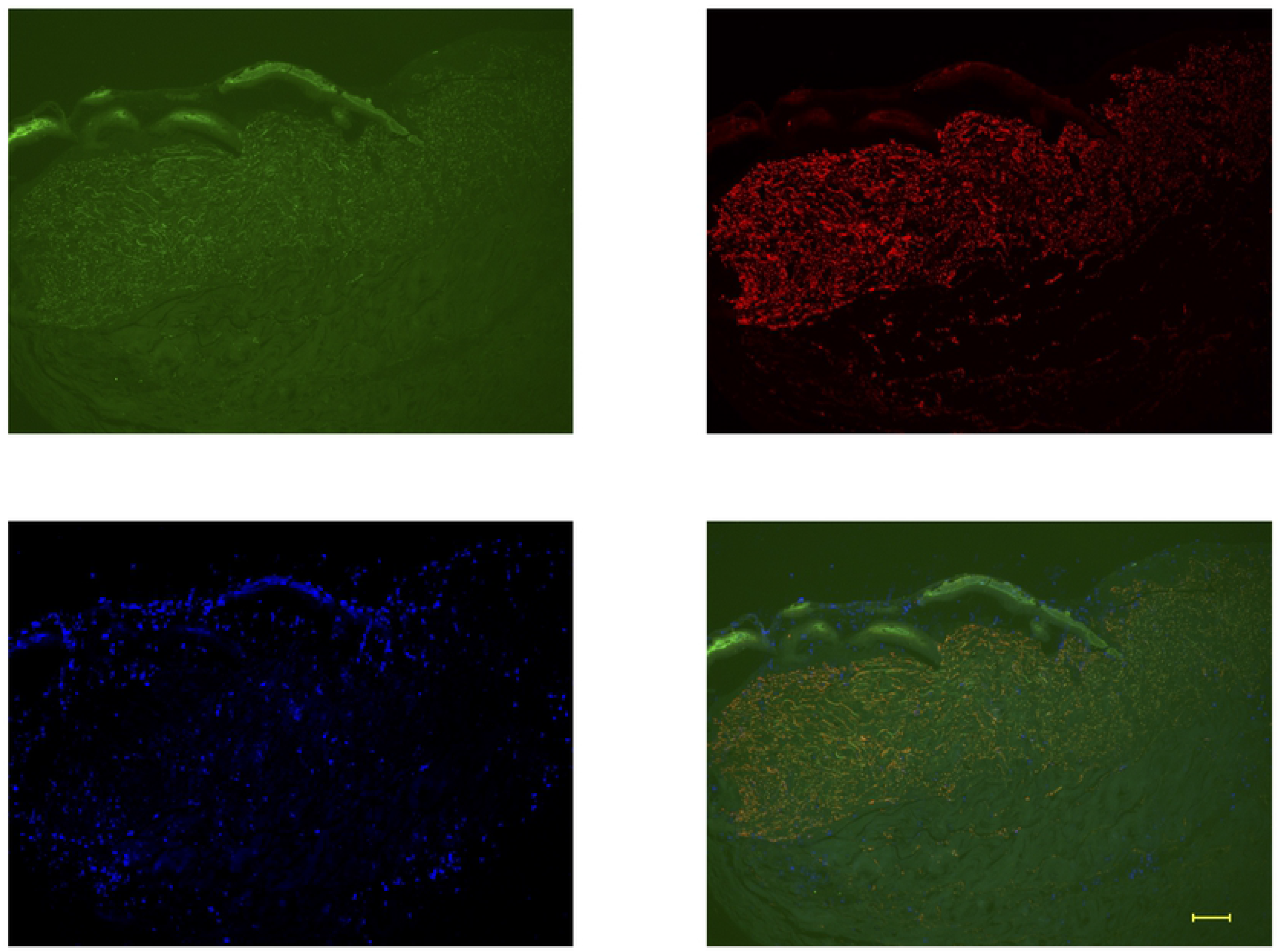
Double fluorescent immunostaining of periosteum at 4 weeks. Scale bar: 100μm. Osteocalcin: green, FBXW2: red, DAPI: blue

**Fig 8.**
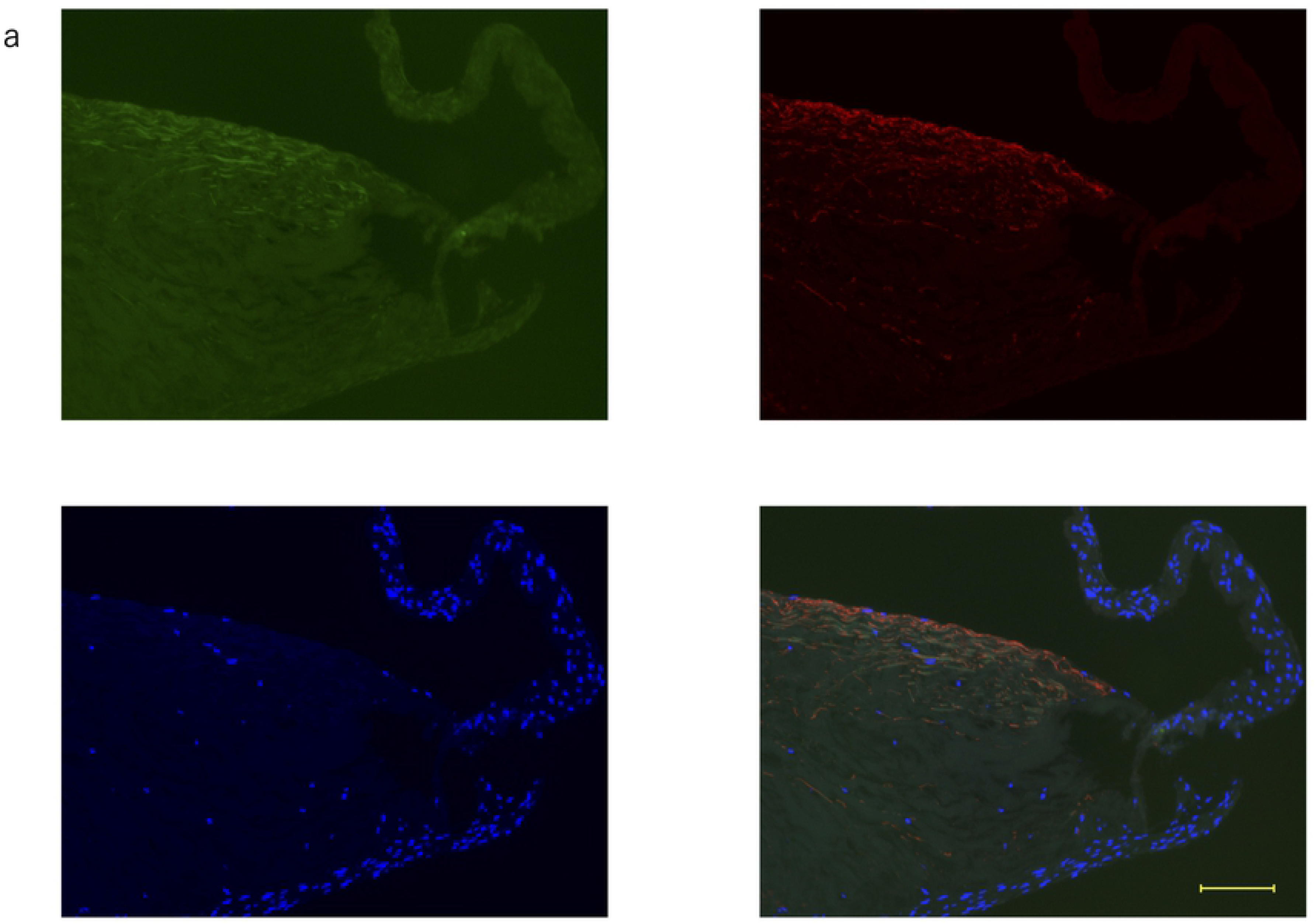
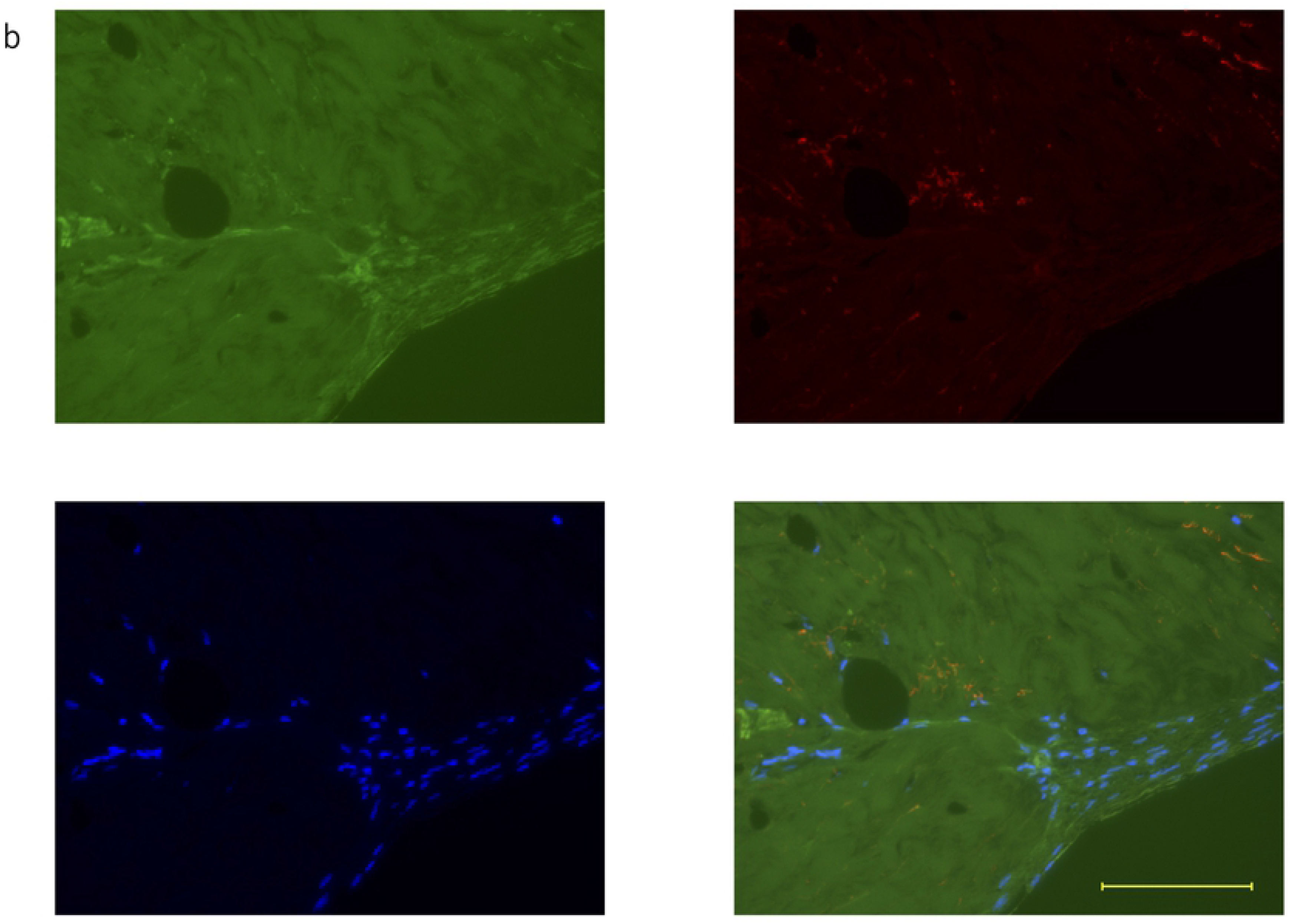
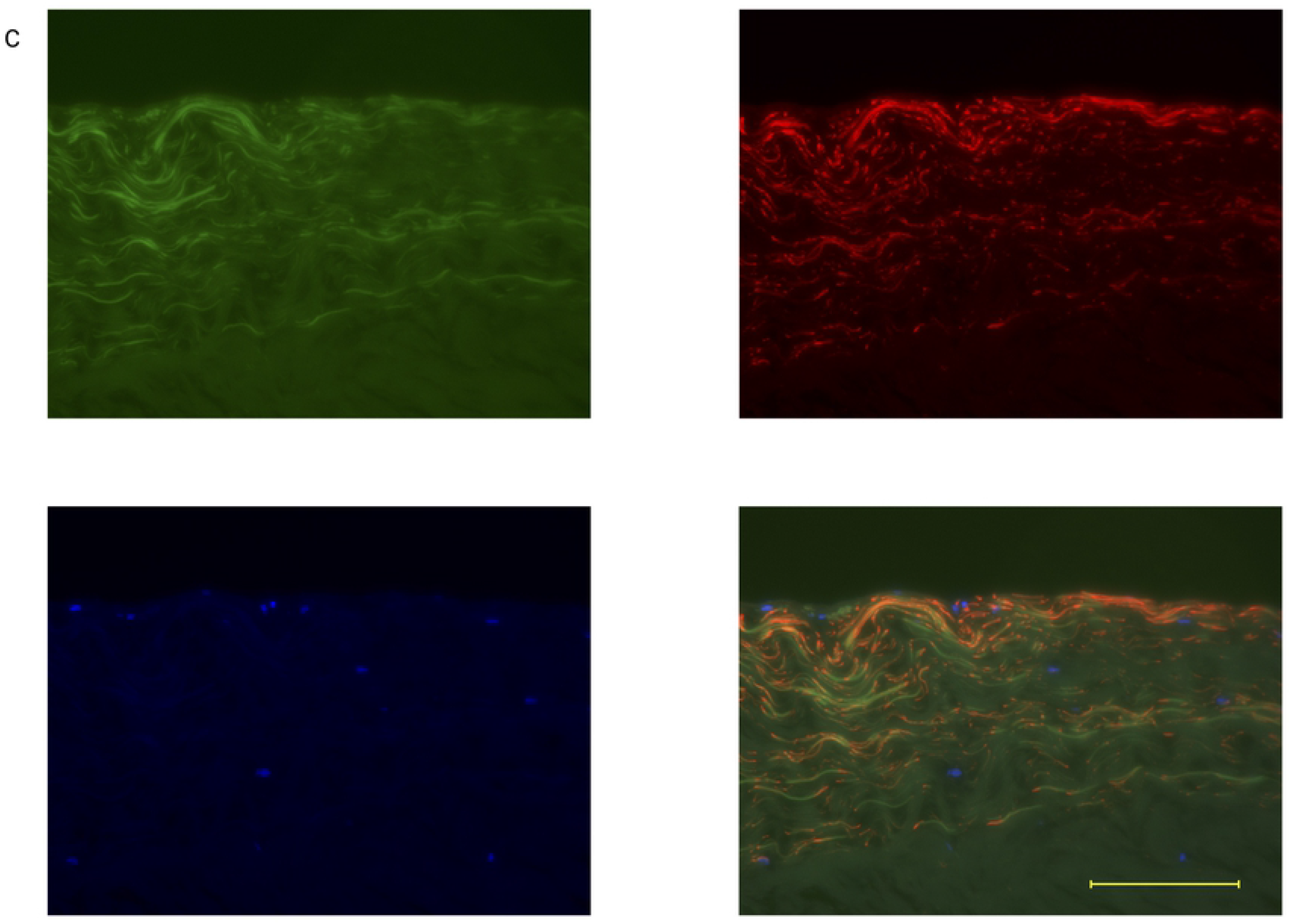
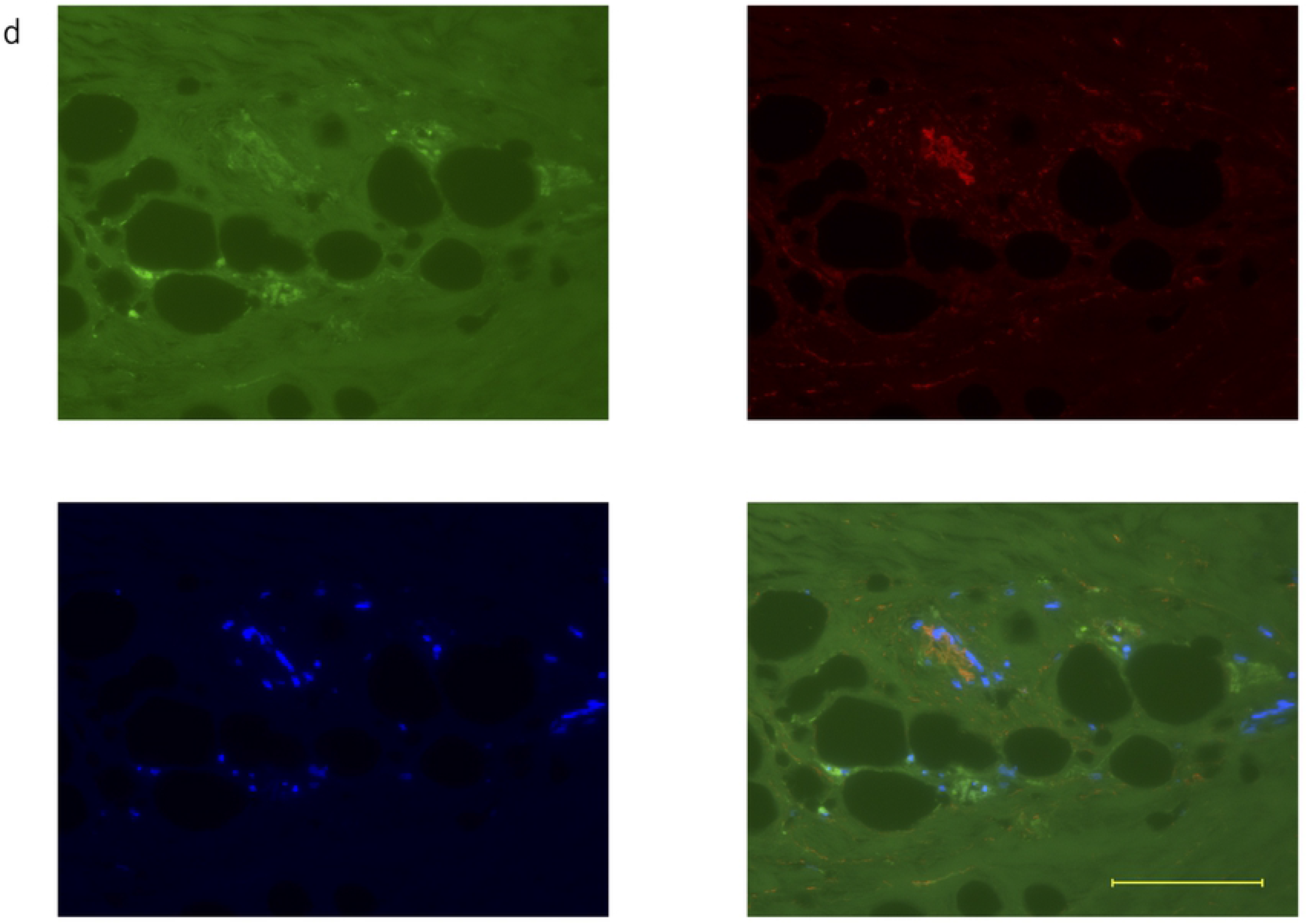
(a)-(d) Double fluorescent immunostaining of periosteum at 5 weeks. Scale bar: 100 μm. Osteocalcin: green, FBXW2: red, DAPI: blue

## Discussion

This study highlights the role of FBXW2. The study revealed that FBXW2 is located in the bone, cambium layer, periosteum, and capillary. Conversely, FBXW2 disappeared around the bulk of osteocalcin and periosteum-derived cells. On day 0, FBXW2 was expressed in the capillary, but osteocalcin was not expressed around the capillary (Fig 3). At day 5, the capillary-like structure collapsed with FBXW2 (Fig 9). Leonard et al. [14] reported that Sirtuin 1 levels decreased in the internal artery of diabetic patients, while osteocalcin levels increased. Therefore, the relationship between FBXW2 and vascular calcification needs to be studied. My observation of FBXW2 expression from day 0 to 5 weeks indicates that FBXW2 plays a role in maintaining tissue shape. It can be hypothesized from Fig 8(a) that FBXW2 maintains the shape of the periosteum and prevents cells from migrating out, and the disappearance of FBXW2 causes migration of periosteum-derived cells out of the periosteal tissue. I further hypothesized that at day 5, the capillary that was no longer needed was disassembled along with FBXW2. In 2015, Hirashima et al. [15] revealed the anchoring structure of the calvarial periosteum. However, the components of the anchoring structure are not clear. In this study, the cambium layer of the periosteum needs FBXW2 to adhere to the bone. Bone is a hard tissue and requires less FBXW2 than the cambium layer. Sun’s group investigated the relationship between FBXW2 and lung cancer cells [16,17]. They reported that FBXW2 suppressed lung cancer cell migration and invasion by inhibiting the escape of these cells.

**Fig 9.**
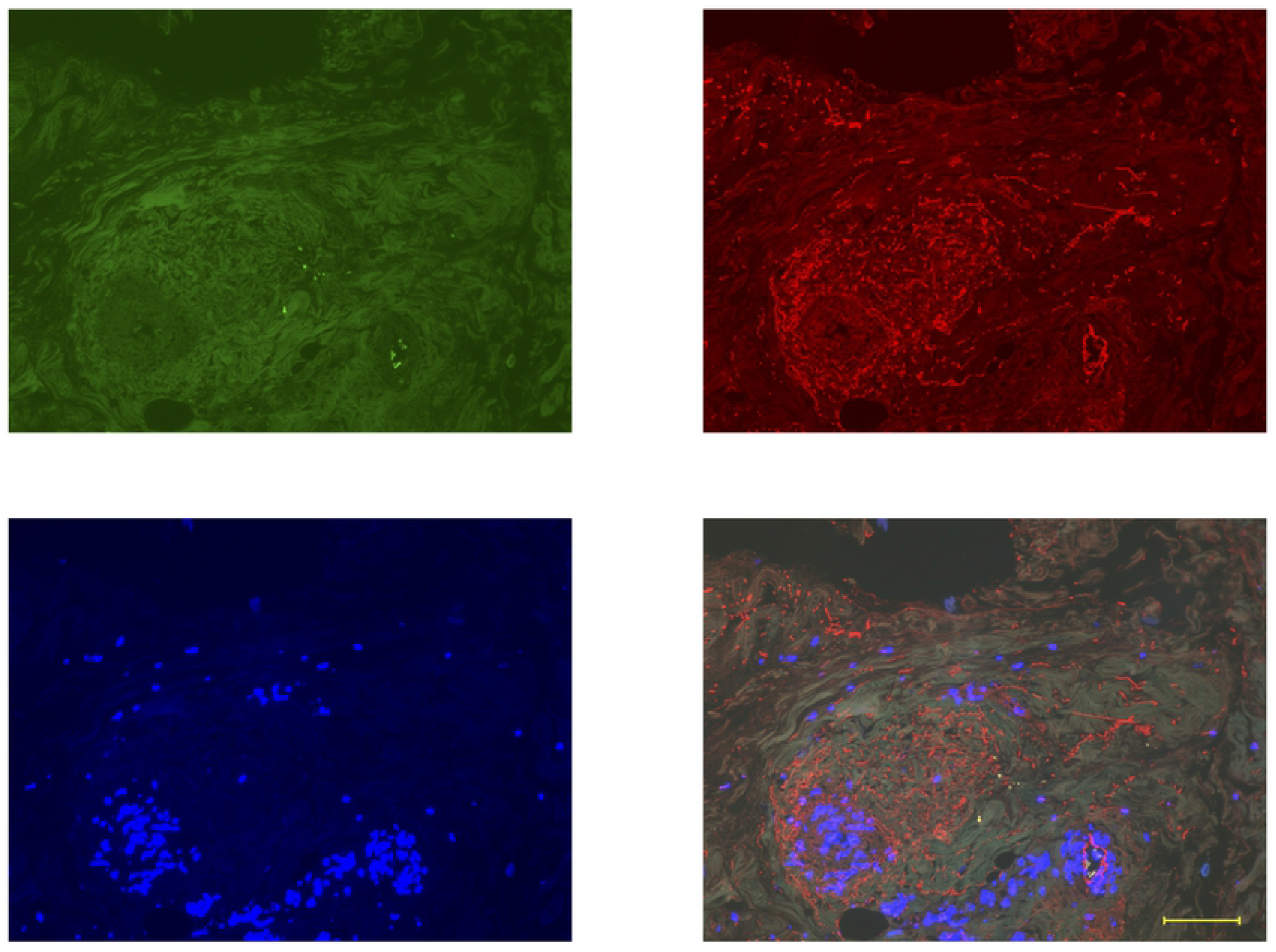
Double fluorescent immunostaining of periosteum at day 5. Scale bar: 100μm. Osteocalcin: green, FBXW2: red, DAPI: blue

The relationship between FBXW2 and osteocalcin has not been established. In 2018, Akiyama [13] reported that FBXW2 was localized with osteocalcin first. Recent studies have reported a relationship between osteocalcin and diabetes [14,18–20]. In this study, osteocalcin was localized in the bone, but RANKL was not. Osteocalcin is an important bone matrix protein [21] and a typical biomarker for bone maturation [22], whereas RANKL is crucial for osteoclastogenesis [23–25]. Thus, in this study, RANKL was not observed as this period did not involve osteoclast formation. Two antibodies for osteocalcin, a mouse monoclonal antibody for osteocalcin (No. sc-376835), and monoclonal antibody to bovine osteocalcin (code no. M042), were used. To compare with the RANKL antibody, no. sc-376835 was used, but for double fluorescent immunostaining, code no. M042 was used to obtain a sharp contrast. No. sc-376835 is of mouse origin, whereas code no. M042 is bovine osteocalcin, the latter being more sensitive. On day zero, osteocalcin was not expressed in the periosteum. At one week, a small amount of osteocalcin appeared with FBXW2 (Fig 4(a)). In a previous study, at five weeks, both FBXW2 and osteocalcin formed long and thin tubes [13]. It is hypothesized that osteocalcin-forming cells have an affinity for FBXW2. If FBXW2 has a role in tissue shape maintenance, it can be assumed that osteocalcin was synthesized on a template of FBXW2. At 2 weeks, the bulk of osteocalcin was separated from FBXW2 with the exception of tube formation (Fig 5(a)–(c)). In Fig 5(f), the antibody for RANKL was used as a negative control under the same conditions (mouse IgG_1_ concentration and treatment time were 4 μg/mL and 4 h, respectively). At three weeks, periosteum-derived cells started to migrate out of the periosteum (Fig 6b) and FBXW2 expression decreased at the root of periosteum-derived cells. At 4 weeks, osteocalcin burst out of the periosteum and fell onto periosteum-derived cells. At 5 weeks, multilayered periosteum-derived cells appeared and FBXW2 disappeared in periosteum-derived cells.

Akiyama et al. [7] reported that transplanted bovine periosteum-derived cells can form new bone. Osteoclasts were followed using TRAP [10] from periosteum-derived cells up to the formation of new bone. Therefore, RANKL, a biomarker of osteoclasts, may be expressed even after transplantation. An important characteristic of periosteum-derived cells is that they can form multilayered cell sheets *in vitro* without artificial scaffold materials [8]. In this study, primary cultured cells were investigated. However, secondary passage cells cannot form multilayered cell sheets (data not shown). In Fig 8(a), periosteum-derived cells seem to form multilayered cell sheets at the inner periosteum and then migrate out of the periosteum. Therefore, secondary passage cells without periosteum cannot form multilayered cell sheets. The important question here is whether a multilayered cell sheet on tissue culture dishes is formed in the periosteum or outside it. Multilayered cell sheets of periosteum-derived cells can carry out scaffold-free bone regeneration [7]. FBXW2 in the periosteum may contribute to multilayered cell sheet formation. Simon et al. [26] reported that surgical stimulation of the periosteum by a sharp incision causes cambium cell proliferation and new bone formation. They concluded that the reason for new bone formation was an increase in the thickness of the cambium layer. However, my results suggest that FBXW2 and osteocalcin *in vitro* are also involved in bone formation *in vivo*. However, I cannot conclude whether FBXW2 and osteocalcin are directly or indirectly related. Other plural proteins may be related to FBXW2 and osteocalcin. FBXW2 is not known to have osteoblastic function. However, osteocalcin is a biomarker of osteoblasts, and signaling pathways for osteogenic differentiation with osteocalcin have been reported [27–32]. In the future, signaling between FBXW2 and osteocalcin should be investigated. Determination of the osteoblastic role of FBXW2 may provide clues for the treatment of osteoporosis.

## Conclusion

1. FBXW2 has a role in maintenance of tissue shape and synthesis of osteocalcin.
2. The disappearance of FBXW2 results in the release of periosteum-derived cells, while osteocalcin remains unaffected.

## Author contribution statement

Mari Akiyama designed and performed the experiments, analyzed and interpreted the data, and wrote the paper.

## Acknowledgements

I thank Kobe Chuo Chikusan for providing the bovine legs, KAC Co., Ltd. for serial sections. I also thank Editage [http://www.editage.com] for editing and reviewing this manuscript for English language. This study was performed at the Institute of Dental Research, Osaka Dental University (Dental Bioscience Facilities).

## References

1. Malden N, Loh S. Pharmacology: Discontinuation of bisphosphonates. Br Dent J. 2017;222(2): 67–68. Epub 2017/01/28. doi: 10.1038/sj.bdj.2017.52. PubMed PMID: 28127004.

2. Roberts SJ, van Gastel N, Carmeliet G, Luyten FP. Uncovering the periosteum for skeletal regeneration: the stem cell that lies beneath. Bone. 2015;70: 10–18. Epub 2014/09/07. doi: 10.1016/j.bone.2014.08.007. PubMed PMID: 25193160.

3. Chang H, Knothe Tate ML. Concise review: the periosteum: tapping into a reservoir of clinically useful progenitor cells. Stem Cells Transl Med. 2012;1(6): 480–491. Epub 2012/12/01. doi: 10.5966/sctm.2011-0056. PubMed PMID: 23197852; PubMed Central PMCID: PMCPMC3659712.

4. Shimizu T, Sasano Y, Nakajo S, Kagayama M, Shimauchi H. Osteoblastic differentiation of periosteum-derived cells is promoted by the physical contact with the bone matrix in vivo. Anat Rec. 2001;264(1): 72–81. Epub 2001/08/16. doi: 10.1002/ar.1126. PubMed PMID: 11505373.

5. Duchamp de Lageneste O, Julien A, Abou-Khalil R, Frangi G, Carvalho C, Cagnard N, et al. Periosteum contains skeletal stem cells with high bone regenerative potential controlled by Periostin. Nat Commun. 2018;9(1): 773. Epub 2018/02/24. doi: 10.1038/s41467-018-03124-z. PubMed PMID: 29472541; PubMed Central PMCID: PMCPMC5823889.

6. Debnath S, Yallowitz AR, McCormick J, Lalani S, Zhang T, Xu R, et al. Discovery of a periosteal stem cell mediating intramembranous bone formation. Nature. 2018;562(7725): 133–139. Epub 2018/09/27. doi: 10.1038/s41586-018-0554-8. PubMed PMID: 30250253; PubMed Central PMCID: PMCPMC6193396.

7. Akiyama M, Nonomura H, Kamil SH, Ignotz RA. Periosteal cell pellet culture system: a new technique for bone engineering. Cell Transplant. 2006;15(6): 521–532. Epub 2006/11/24. doi: 10.3727/000000006783981765. PubMed PMID: 17121163.

8. Akiyama M, Nakamura M. Bone regeneration and neovascularization processes in a pellet culture system for periosteal cells. Cell Transplant. 2009;18(4): 443–452. Epub 2009/07/23. doi: 10.3727/096368909788809820. PubMed PMID: 19622231.

9. Akiyama M. Identification of UACA, EXOSC9, and ΤΜX2 in bovine periosteal cells by mass spectrometry and immunohistochemistry. Anal Bioanal Chem. 2014;406(24): 5805–5813. Epub 2014/04/04. doi: 10.1007/s00216-014-7673-3. PubMed PMID: 24696107.

10. Akiyama M. Characterization of the F-box Proteins FBXW2 and FBXL14 in the Initiation of Bone Regeneration in Transplants given to Nude Mice. Open Biomed Eng J. 2018;12: 75–89. Epub 2018/11/20. doi: 10.2174/1874120701812010075. PubMed PMID: 30450135; PubMed Central PMCID: PMCPMC6198513.

11. Ho MS, Tsai PI, Chien CT. F-box proteins: the key to protein degradation. J Biomed Sci. 2006;13(2): 181–191. Epub 2006/02/08. doi: 10.1007/s11373-005-9058-2. PubMed PMID: 16463014.

12. Wang Z, Liu P, Inuzuka H, Wei W. Roles of F-box proteins in cancer. Nat Rev Cancer. 2014;14(4): 233–247. Epub 2014/03/25. doi: 10.1038/nrc3700. PubMed PMID: 24658274; PubMed Central PMCID: PMCPMC4306233.

13. Akiyama M. FBXW2 localizes with osteocalcin in bovine periosteum on culture dishes as visualized by double immunostaining. Heliyon. 2018;4(9): e00782. Epub 2018/09/20. doi: 10.1016/j.heliyon.2018.e00782. PubMed PMID: 30229138; PubMed Central PMCID: PMCPMC6141272.

14. Bartoli-Leonard F, Wilkinson FL, Schiro A, Inglott FS, Alexander MY, Weston R. Suppression of SIRT1 in diabetic conditions induces osteogenic differentiation of human vascular smooth muscle cells via RUNX2 signalling. Sci Rep. 2019;9(1): 878. Epub 2019/01/31. doi: 10.1038/s41598-018-37027-2. PubMed PMID: 30696833; PubMed Central PMCID: PMCPMC6351547.

15. Hirashima S, Ohta K, Kanazawa T, Uemura K, Togo A, Yoshitomi M, et al. Anchoring structure of the calvarial periosteum revealed by focused ion beam/scanning electron microscope tomography. Sci Rep. 2015;5: 17511. Epub 2015/12/03. doi: 10.1038/srep17511. PubMed PMID: 26627533; PubMed Central PMCID: PMCPMC4667224.

16. Xu J, Zhou W, Yang F, Chen G, Li H, Zhao Y, et al. The β-TrCP-FBXW2-SKP2 axis regulates lung cancer cell growth with FBXW2 acting as a tumour suppressor. Nat Commun. 2017;8: 14002. Epub 2017/01/17. doi: 10.1038/ncomms14002. PubMed PMID: 28090088; PubMed Central PMCID: PMCPMC5241824.

17. Yang F, Xu J, Li H, Tan M, Xiong X, Sun Y. FBXW2 suppresses migration and invasion of lung cancer cells via promoting β-catenin ubiquitylation and degradation. Nat Commun. 2019;10(1): 1382. Epub 2019/03/29. doi: 10.1038/s41467-019-09289-5. PubMed PMID: 30918250; PubMed Central PMCID: PMCPMC6437151.

18. Liu DM, Mosialou I, Liu JM. Bone: Another potential target to treat, prevent and predict diabetes. Diabetes Obes Metab. 2018;20(8): 1817–1828. Epub 2018/04/25. doi: 10.1111/dom.13330. PubMed PMID: 29687585.

19. Zoch ML, Clemens TL, Riddle RC. New insights into the biology of osteocalcin. Bone. 2016;82: 42–49. Epub 2015/06/10. doi: 10.1016/j.bone.2015.05.046. PubMed PMID: 26055108; PubMed Central PMCID: PMCPMC4670816.

20. Brennan-Speranza TC, Conigrave AD. Osteocalcin: an osteoblast-derived polypeptide hormone that modulates whole body energy metabolism. Calcif Tissue Int. 2015;96(1): 1–10. Epub 2014/11/25. doi: 10.1007/s00223-014-9931-y. PubMed PMID: 25416346.

21. Bailey S, Karsenty G, Gundberg C, Vashishth D. Osteocalcin and osteopontin influence bone morphology and mechanical properties. Ann N Y Acad Sci. 2017;1409(1): 79–84. Epub 2017/10/19. doi: 10.1111/nyas.13470. PubMed PMID: 29044594; PubMed Central PMCID: PMCPMC5730490.

22. Ho MH, Yao CJ, Liao MH, Lin PI, Liu SH, Chen RM. Chitosan nanofiber scaffold improves bone healing via stimulating trabecular bone production due to upregulation of the Runx2/osteocalcin/alkaline phosphatase signaling pathway. Int J Nanomedicine. 2015;10: 5941–5954. Epub 2015/10/10. doi: 10.2147/ijn.S90669. PubMed PMID: 26451104; PubMed Central PMCID: PMCPMC4590342.

23. Kartsogiannis V, Zhou H, Horwood NJ, Thomas RJ, Hards DK, Quinn JM, et al. Localization of RANKL (receptor activator of NF kappa B ligand) mRNA and protein in skeletal and extraskeletal tissues. Bone. 1999;25(5): 525–534. Epub 1999/11/26. doi: 10.1016/s8756-3282(99)00214-8. PubMed PMID: 10574572.

24. Ikebuchi Y, Aoki S, Honma M, Hayashi M, Sugamori Y, Khan M, et al. Coupling of bone resorption and formation by RANKL reverse signalling. Nature. 2018;561(7722): 195–200. Epub 2018/09/07. doi: 10.1038/s41586-018-0482-7. PubMed PMID: 30185903.

25. Siddiqui JA, Partridge NC. Physiological bone remodeling: Systemic Regulation and growth factor involvement. Physiology (Bethesda). 2016;31(3): 233–245. Epub 2016/04/08. doi: 10.1152/physiol.00061.2014. PubMed PMID: 27053737; PubMed Central PMCID: PMCPMC6734079.

26. Simon TM, Van Sickle DC, Kunishima DH, Jackson DW. Cambium cell stimulation from surgical release of the periosteum. J Orthop Res. 2003;21(3): 470–480. Epub 2003/04/23. doi: 10.1016/s0736-0266(02)00206-1. PubMed PMID: 12706020.

27. García-Gareta E, Coathup MJ, Blunn GW. Osteoinduction of bone grafting materials for bone repair and regeneration. Bone. 2015;81: 112–121. Epub 2015/07/15. doi: 10.1016/j.bone.2015.07.007. PubMed PMID: 26163110.

28. Hassumi JS, Mulinari-Santos G, Fabris A, Jacob RGM, Gonçalves A, Rossi AC, et al. Alveolar bone healing in rats: micro-CT, immunohistochemical and molecular analysis. J Appl Oral Sci. 2018;26: e20170326. Epub 2018/06/14. doi: 10.1590/1678-7757-2017-0326. PubMed PMID: 29898174; PubMed Central PMCID: PMCPMC6010327.

29. Miron RJ, Sculean A, Shuang Y, Bosshardt DD, Gruber R, Buser D, et al. Osteoinductive potential of a novel biphasic calcium phosphate bone graft in comparison with autographs, xenografts, and DFDBA. Clin Oral Implants Res. 2016;27(6): 668–675. Epub 2015/08/01. doi: 10.1111/clr.12647. PubMed PMID: 26227281.

30. Oh SH, Kim JW, Kim Y, Lee MN, Kook MS, Choi EY, et al. The extracellular matrix protein Edil3 stimulates osteoblast differentiation through the integrin α5β1/ERK/Runx2 pathway. PLoS One. 2017;12(11): e0188749. Epub 2017/11/29. doi: 10.1371/journal.pone.0188749. PubMed PMID: 29182679; PubMed Central PMCID: PMCPMC5705136.

31. Tang Z, Li X, Tan Y, Fan H, Zhang X. The material and biological characteristics of osteoinductive calcium phosphate ceramics. Regen Biomater. 2018;5(1): 43–59. Epub 2018/02/10. doi: 10.1093/rb/rbx024. PubMed PMID: 29423267; PubMed Central PMCID: PMCPMC5798025.

32. Avery SJ, Sadaghiani L, Sloan AJ, Waddington RJ. Analysing the bioactive makeup of demineralised dentine matrix on bone marrow mesenchymal stem cells for enhanced bone repair. Eur Cell Mater. 2017;34: 1–14. Epub 2017/07/12. doi: 10.22203/eCM.v034a01. PubMed PMID: 28692113.

